# SB 11285, a Novel Canonical Purine-Pyrimidine Cyclic Dinucleotide, is a Systemically Bioavailable STING Agonist for Cancer Immunotherapy

**DOI:** 10.1101/2025.03.27.645620

**Authors:** Sreerupa Challa, Shenghua Zhou, Anjaneyulu Sheri, Geeta Meher, Seetharamaiyer Padmanabhan, Santosh Khedkar, Vishal Nair, Rayomand Gimi, Dillon Cleary, Radhakrishnan P. Iyer

**Author notes:** These authors contributed equally to the project (Sreerupa Challa, Shenghua Zhou).

## Abstract

**Summary:** SB 11285, a novel cell-permeable, canonical 3’,5, purine-pyrimidine cyclic dinucleotide has been developed as a potent systemically bioavailable STING agonist currently in clinical trials.

The stimulator of Interferon genes (STING) pathway, crucial for cytosolic DNA sensing and innate immunity, presents a promising avenue for novel cancer immunotherapies. Discussed here is the discovery and development of SB11285, a potent activator of the STING pathway, belonging to the canonical class of 3’,5’ purine-pyrimidine (PuPy) cyclic dinucleotides (CDNs). SB11285 exhibits superior potency and cell permeability compared to non-canonical 2,3’-CDNs and demonstrates robust antitumor activity in in vivo syngeneic mouse tumor models, resulting in complete tumor regression, and durable immunological memory. Combination with checkpoint inhibitors and cytotoxic agents further enhances its efficacy. The anti-tumor effects of SB 11285 rely on the induction of CD8+ T cells, activated macrophages, and NK cells in the tumor microenvironment (TME), for efficient tumor killing and fostering profound and durable anti-tumor response. Our findings underscore the potential of functionalized canonical 3’,5’-Pu-Py CDNs, exemplified by SB11285, as safe and potent STING agonists for multiple routes of administration in cancer therapy. Currently, SB11285 administered intravenously (IV), is undergoing evaluation as the first systemically administered CDN analog in human clinical trials across various tumor types.

## Introduction

Cancer immunotherapy, which harnesses the body’s immune system for anti-tumor response, is now a front-line therapeutic approach, especially for cancers resistant to other treatments. Recent studies indicate that many patients with advanced solid tumors exhibit a phenotype of a spontaneous T cell inflamed TME, which correlates with favorable clinical responses to immunotherapies (1–9). The inflamed TME is characterized by a type I interferon (IFN) signature that is associated with T cell activation against tumor antigens and the generation of CD8+ T cells for tumor eradication (9). The STING pathway, responsible for cytosolic DNA sensing, is recognized as a critical innate immune mechanism driving type I IFN production (9–12). Consequently, the discovery of compounds activating the STING pathway has become a promising area for developing novel immunotherapeutic strategies against cancer.

STING (also known as TMEM173, MITA, ERIS, and MPYS) is a cellular adapter protein activated by cyclic dinucleotides (CDNs), which are synthesized by the cyclic GMP-AMP synthase (cGAS) enzyme in response to cytosolic DNA from tumors or viral infections (10–12). The STING protein is composed of a homodimer with 379 amino acid (aa) residues, featuring an *N*-terminal multi-putative transmembrane domain located in the endoplasmic reticulum (ER) (aa regions 1–154), a dimerization and ligand-binding domain (aa 155–342), and a *C*-terminal tail (CTT, aa 343–379) receptive to phosphorylation by Tank-binding kinase 1 (TBK1) (10–12). Upon binding of ligands such as CDNs, STING undergoes activation, conformational changes, and translocation from the ER, through the Golgi, to a distinct perinuclear region where it forms aggregates (11,12). TBK1 also traffics in a STING-dependent manner and phosphorylates STING, subsequently activating the transcription factors such as IRF3 and IRF7 that leads to type I interferon (IFN-α and IFN-β) production thereby stimulating the immune system. Besides sensing tumor-derived DNA, this pathway is implicated in DNA virus sensing as well (12). Aberrant STING activation is associated with multiple autoimmune and inflammatory diseases (13–15). Furthermore, innate immune sensing in the TME is crucial for promoting tumor-initiated T cell priming and induction of tumor-infiltrating lymphocytes (TILs) for antitumor activity (16).

IFN-β serves as the hallmark cytokine triggered upon STING activation, either by exogenous CDNs produced during bacterial infection or by a distinct endogenous CDN synthesized by host cyclic GMP-AMP synthetase (cGAS) upon the sensing of cytosolic double-stranded DNA (dsDNA) (17–23). Canonical 3’,5’-linked CDNs like c-di-GMP and c-di-AMP, derived from guanosine and adenosine, are common in bacteria, archaea, and protozoa (20–22). In contrast, following DNA transfection or DNA virus infection in mammalian cells, cGAS generates a unique non-canonical CDN, 2’,3’-cGAMP (cGAMP), featuring guanine and adenine as purine nucleobases. cGAMP, the first CDN found in metazoans through cGAS sensing of cytosolic DNA, activates STING to induce the production of type I IFNs (24–27). With a distinctive 2’,5’ internucleotidic phosphodiester bond, different from the canonical 3’,5’-linked bacterial CDNs, cGAMP is hypothesized to exhibit unique signaling properties, including resistance to endogenous 3’,5’ phosphodiesterase and the capability to bind multiple allelic variants of STING present in the human population (25–27). Consequently, several 2’,5’-CDNs, with both nucleobases being purines, have emerged as potent vaccine adjuvants (28–33), and CDNs such as ADU-S100 and MK-1454 have progressed towards human clinical trials as immunotherapeutic agents against cancer (34–35). However, the efficacy studies of ADU-S100 in syngeneic mouse tumor models indicated the necessity for intratumoral (IT) injection to achieve maximal therapeutic effect, potentially limiting its clinical utility to tumors accessible via IT administration (36).

Previously, we reported that the dinucleotide SB 9200 acts as an agonist of retinoic acid-inducible gene 1 (RIG-I), exhibiting broad-spectrum antiviral activity against HBV, HCV, RSV, and influenza virus, (37,38) as well as, adjuvant activity when combined with the Mycobacterium tuberculosis (MTb) vaccine (39). SB 9200 is a linear 3’,5’-linked dinucleotide composed of adenosine and uridine, which are purine, and pyrimidine nucleotides, respectively. Intrigued by this finding, we explored the potential of the corresponding PuPy-CDNs as STING agonists. Extensive structure-activity relationship (SAR) studies of PuPy-CDNs involving the modifications of sugar, nucleobase, and phosphodiester groups, as well as variations in internucleotidic connectivities, were carried out that resulted in a library of several hundred PuPy-CDNs. These CDNs were evaluated for STING-dependent IFN-induction using cell-based reporter assays. Remarkably, we discovered that chemo-selective attachment of ar-alkyl chains of varying lengths to the lone phosphorothioate backbone of a PuPy-CDN significantly enhanced STING-dependent IFN and NF-KB induction (40). Moreover, unlike ADU-S100 and cGAMP, some of these novel CDNs, including the lead compound SB 11285, a 3’,5’-PuPy-CDN, were taken up by cells without transfection reagents, and could activate multiple allelic variants of human STING (hSTING). Further investigations using human peripheral blood mononuclear cells (PBMCs) revealed that SB 11285 was preferentially taken up by dendritic cells (DCs) and monocytes.

We performed computational modeling and molecular dynamic simulations of PuPy-CDNs with STING using the published crystal structures of c-di-GMP or cGAMP with STING protein (25–27). Our analysis revealed that in addition to the Pu-Py interactions with specific amino acid residues of STING, the ar-alkyl chain of SB 11285 interacted with amino acid residues in the C-terminal domain (CTD) (amino acid residues 139-344) of STING, thereby significantly enhancing the binding of SB 11285 to STING. The computational observations were further supported by binding studies of SB 11285 using surface plasmon resonance with STING-CTD containing a C-terminal tail (CTT, aa residues 337–379). In pharmacology studies, the administration of SB 11285 via intratumoral (IT), intravenous (IV), or intraperitoneal (IP) routes demonstrated potent anti-tumor activity in multiple subcutaneous and orthotopic syngeneic mouse and rat tumor models, that was additionally associated with the induction of immune memory and abscopal effects. Pharmacodynamic studies in syngeneic mouse tumor models demonstrated the production of type I IFNs and multiple antitumor cytokines, along with the induction of CD8+ T cells and reduction of regulatory T cells (Tregs). These studies were supportive of the activation of STING pathway by SB 11285. Our preclinical studies highlight the high translational potential of SB 11285 as an immunotherapeutic agent against cancer. Moreover, our findings suggest that an expanded repertoire of cell-permeable 3’,5’, 2’,3’, and 3’,2’-PuPy-CDN compounds can be developed as STING agonists for therapeutic applications, including antivirals, antitumor agents, and vaccine adjuvants, that can be administered via multiple routes.

## Materials and Methods

### Synthesis of Compounds

The library of PuPy-CDNs were synthesized and purified (>95% purity) by high-performance liquid chromatography (HPLC) according to reported procedures (40). The lead compounds were purified to > 98% purity and were characterized by ^1^H NMR, ^31^P NMR, LC-MS, and HPLC and certified free from endotoxin before use in preclinical studies. Biotin-tagged compounds were synthesized by standard phosphoramidite chemistry (41) using the appropriate biotin building block (Glen Research).

### Cells

HEK293, HEK-Blue ISG, HEK-Blue ISG-KO-STING, 293/hTLR7-HA, 293/hTLR9-HA, THP1-Blue ISG, THP1-Dual, THP1-Dual-KO-STING, THP1-Dual-MyD, as well as, RAW-Lucia ISG, RAW-Lucia-KO-STING, RAW-Lucia-KO-TBK1, RAW-Lucia-KO-IRF3, RAW-Lucia-KO-IRF7, RAW-Lucia-KO-MAVS, RAW-Lucia-KO-TRIF. HEK293T and NK-92 cell lines were purchased from InvivoGen and ATCC. HEK293, and HEK293T cells were maintained in complete DMEM supplemented with 10% heat-inactivated FBS, 100 U/mL penicillin, 100 µg/mL streptomycin, 2 µM L-glutamine. THP1, NK-92 and RAW cells THP1, NK-92 and RAW cells were maintained in complete RPMI-1640 supplemented with 10% heat-inactivated FBS, 100 U/mL penicillin, 100 µg/mL streptomycin, 2 µM L-glutamine. SZ14 ISRE-luc reporter cell line is an in-house generated HEK293-derived stably expressing ISG54 ISRE-luc and Renilla-luc (Stratagene) and maintained in complete DMEM supplemented with 10% heat-inactivated FBS, 100 U/mL penicillin, 100 µg/mL streptomycin, 100 µg/mL hygromycin, 2 µM L-glutamine.

Primary human healthy PBMCs (fresh and frozen) were purchased from ZenBio, Inc. Primary mouse wild-type and STING-KO splenocytes and bone marrow-derived macrophages were prepared from wild-type and STING-KO mice (purchased from the Jackson Laboratory).

Scanning Electron Microscopy (SEM), Imaging and Immunohistochemistry (IHC) of tissue samples were performed using the Core facilities at the University of Massachusetts Medical Center, Worcester, MA.

### Plasmids

The plasmids pUNO1-hSTING-HA3x, and pUNO1-hSTING-MRP were purchased from InvivoGen. The pISRE-luc and pNF-κB-Luc cis-reporter plasmids were purchased from Stratagene. pRL-TK plasmid was obtained from Promega.

The following STING mutants were generated from the parental plasmid pUNO1-hSTING3x using QuickChange II site-directed mutagenesis kit (Agilent): STING-R232, -Y167A, -R238A, -Y240A, -N242A, -E260A, -T263A, -R232A, -S366A. Mutations were verified by DNA sequencing (Genewiz). Plasmids were purified using HiSpeed plasmid midi-, and maxi kits (Qiagen). Plasmids were transfected using GeneJuice (Sigma) by following the manufacturer’s recommendation.

### STING CTD with CTT Expression

Expression and purification of purified human STING C-terminal domain protein with C-terminal tail CTT (amino acids 139 to 379) and C-terminal domain without C-terminal tail CTT (amino acids 139 to 341) were performed. Briefly, the 6 x His STING CTD open reading frame was cloned into piTX-1 gene (Invitrogen) and used to transform the Escherichia coli strain pLysS (Promega). The transformed *E. coli* cells were then induced to express the STING protein using 1 mM isopropyl-D-thiogalacto-pyranoside (IPTG) at 16°C for 18 hrs. STING protein was purified by nickel-affinity chromatography and further purified by gel filtration chromatography (HiPrep 16/60 Sephacryl S-100 HR column; GE Healthcare Life Sciences). Eluted proteins were concentrated using Amicon centrifugal filters (3 kDa cutoff).

### In Vitro Screening

HEK293 cells were co-transfected with expression plasmids for the ISG54 ISRE-firefly luciferase reporter gene (ISRE-luc) (Stratagene) and the hygromycin resistance gene (InvivoGen) using GeneJuice transfection reagent followed by exposure to hygromycin to select SZ14 as the best representative stable clone, robustly responding to treatment with Sendai virus (ATCC) and 2’,3’-cGAMP (InvivoGen) (data not shown).

Primary screening was carried out in a 96-well plate format. SZ14 cells were seeded into white 96-well Lumitrac tissue culture polystyrene plates (Promega) in triplicate at 10^5^/150 µl/well and incubated for 2 hr., at 37 °C, in 5% CO_2_ humidified incubator. Cells were treated with a single concentration at 10 μM of each 3’,5’-PuPy-CDN or DMSO vehicle control, or 10 μM cGAMP at 50 µl/well in the presence of digitonin (10 µg/mL) for 30 min. The medium was then aspirated and fresh DMEM supplemented with 10% heat inactivated FBS, Pen/Strep, and L-glutamine was added at 100 µl/well. After incubation for 5 hr. at 37 °C, medium was removed, 35 µL/well of 1:1 pre-diluted/DPBS buffered Steady-Glo luciferase buffer (Promega) was added, and luciferase activity read on Infinite Pro200 plate reader and plotted as fold-increase over ISRE-luc activity in DMSO-treated cells. Positive “hits” were identified that showed > 5-fold increase in ISRE-luc activity as compared to DMSO-treated cells. EC_50_ and CC_50_ values of lead compounds were calculated using XLfit.

### Kinetics of signaling by activated STING

SB 11285 activation is characterized by signaling proteins as indicated by the formation of phosphorylated IRF3, TBK-1 and IκB-α, detectable after either SB 11285 or cGAMP treatment. THP-1 cells were treated with either 5 µM SB 11285 or cGAMP with Lipofectamine LTX, and at different time-points, whole cell extracts were treated with RIPA lysis buffer (Promega) and analyzed by immunoblotting with anti-hSTING antibody, anti-IRF3, anti-phospho-IRF3, anti-TBK1, anti-phospho TBK1, anti-IKBα, anti-phospho IKBα, β-actin antibodies and anti-rabbit IgG, HRP-linked antibody.

### Activity of SB 11285 Against STING Allelic Variants

To evaluate whether SB 11285 activates various STING allelic variants, THP1 cells that naturally express R71H, G230A and R293Q (HAQ) HAQ isoform and STING-KO THP1 cells, stably transfected with WT or allelic variants, and carrying both an ISRE-inducible Lucia and an NF-κB inducible SEAP reporters, were treated with various concentrations of SB 11285 or DMSO control. After 20 hrs., the levels of Lucia and SEAP were measured using QUANTI-luc and QUANTI-Blue to measure ISRE and NF-κB reporter activity respectively. The % IRF Induction was calculated from fold-change in luminescence/absorbance compared to DMSO treated sample. EC_50_ values were generated by curve fit in Xlfit. We also performed similar assays of SB 11285 in HEK293 cells transiently transfected with various in-house generated STING mutants plus ISG54 ISRE-luc and Renilla-luc reporter genes.

### Gene Expression Analysis in THP1, PBMCs and A20 lymphoma Cells

Cells were grown in complete media and were treated with 25 μM SB 11285 or cGAMP or DMSO with Lipofectamine LTX. After 48 hrs. incubation, RNA was extracted, and gene expression was evaluated by Real time PCR. Fold-induction of genes was assessed by calculating the relative fold gene expression of samples by the ΔΔct method.

### In Vivo Tumor Experiments

The test compounds were administered by intratumoral (I.T.), intraperitoneal (I.P.), and intravenous, (I.V.), routes. All in vivo studies were conducted by Charles River Discovery Services, USA, Syngene International, India, and The Jackson Laboratory, USA.

### Study Materials and Preparation of Test articles for In Vivo Studies

SB 11285, and related analogs of >98% purity were certified free from endotoxin (<2.5 EU/mg) (Cape Cod Laboratories, MA, USA). Cyclophosphamide and Docetaxel were of the highest purity that was available from reputed vendors. Anti-PD1 (RMP1-14) and anti-CTLA-4 (9D9) antibodies were purchased from BioXell, NH, USA. Rat IgG2b isotype control, anti-keyhole limpet hemocyanin, and anti-mouse CD8α were obtained from BioXell, NH USA.

SB 11285, and analogs were dissolved by adding sterile saline into pre-weighed compounds in polypropylene tubes, incubating at 37 °C for 5 minutes, followed by sonication. Cyclophosphamide was dissolved in saline and administered in a dosing volume of 10 mL/kg. Docetaxel (stock 40 mg/mL) diluted with diluent and 10% ethanol to a working solution of 1.5 mg/mL and administered at 5 mL/kg. Anti-PD-1 antibody, anti-CTLA antibody, anti-CD8b antibody and rat IgG2b were prepared in PBS and administered at 0.2 mL/mouse.

### Animals

Female BALB/c mice (BALB/c AnNcr1, Charles River), eight to 10 weeks old, on Day 1 of the study with a body weight range of 15 to 21 g, were used for the anti-tumor studies. The animals were provided water *ad libitum* (reverse osmosis, 1 ppm Cl) and NIH 31 Modified and Irradiated Lab Diet® consisting of 18.0% crude protein, 5.0% crude fat, and 5.0% crude fiber. Wistar IGS rats 7-8 weeks old, weighing 180 −220 g were obtained from Vivo Bio Tech. The animal care and use were as per guidelines from the Association for Assessment and Accreditation of Laboratory Animal Care International (AAALAC), which assures compliance with accepted standards for the care and use of laboratory animals.

### Tumor Cell Culture

A20 murine B cell lymphoma cells, CT26 murine colon carcinoma cells, and NBT-II (Nara Bladder Tumor No.2) bladder cancer cells were obtained from the American Type Culture Collection (ATCC). The 4T1-luc2 murine mammary gland tumor cell line was obtained from Perkin-Elmer. MC-38 colon adenocarcinoma cells were obtained from Creative Bioarray.

The tumor cells were maintained as exponentially growing suspension cultures in RPMI-1640 medium supplemented with 10% fetal bovine serum, 4.5 g/L glucose, 2 mM glutamine, 10 mM HEPES, 1 mM sodium pyruvate, 0.075% sodium bicarbonate, 100 units/mL penicillin G sodium, 100 μg/mL streptomycin sulfate, 25 μg/mL gentamicin, and 50 μM β-mercapto-ethanol. The tumor cells were grown in tissue culture flasks in a humidified incubator at 37 °C, in an atmosphere of 5% CO_2_.

### In Vivo Implantation and Tumor Growth in Syngeneic Models

The tumor cells used for implantation were harvested during log-phase growth and re-suspended at a concentration of 1 x 10^5^ to 1 x 10^7^ cells/mL in cold phosphate buffered saline (PBS). Mice were injected subcutaneously in the right flank tumor cells as a 0.1 mL cell suspension in PBS. Fourteen days after tumor cell implantation, animals were sorted into treatment groups (n = 10/group) with individual tumor volumes of 80 to 140 mm^3^. Tumors were measured with a caliper twice weekly for the duration of the study. Separate groups of animals (n = 4/group) were designated for sample collection.

### Tumor Measurements, Tumor Growth Inhibition (TGI) and Tumor Growth Delay (TGD)

Tumors were measured in two dimensions using calipers, and volume was calculated using the formula: Tumor Volume (mm^3^) = L X W^2^ /2 where, L = Length (mm); W = Width (mm) of tumor.

The study endpoint was defined as a mean tumor volume (MTV) of 1500 mm^3^ in the control group or Day 45, whichever came first. When the TGI endpoint was reached, the experiment was converted to a TGD study. The TGD study endpoint was defined as a tumor volume of 2000 mm^3^ or Day 45, whichever came first. Treatment outcome was evaluated from TGD, and was defined as the increase in the median time to end point (TTE) for a treatment group compared to the control group expressed in days: TGD = T – C; or as a percentage of the median TTE of the control group:

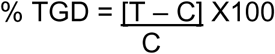

T = median TTE for a treatment group

C = median TTE for the control group

### Statistical and Graphical Analyses

Prism (GraphPad) for Windows 7.02 was used for graphical presentations and statistical analyses. Groups of more than two were analyzed by on-way ANOVA followed by Tukey’s Multiple Comparison Test. Survival was analyzed by the Kaplan-Meier method. Test results are reported as not significant (ns) at *P* > 0.05, significant symbolized by “*”) at 0.01 < *P* ≤ 0.05, very significant (“**”) at 0.001 < *P* ≤ 0.01, and extremely significant (“***”) at *P* ≤ 0.001.

### Sampling for Analysis

The whole blood was collected by terminal cardiac puncture under isoflurane anesthesia into tubes containing K_2_EDTA anti-coagulant and placed on wet ice. Blood (100 μL) was processed for serum in the absence of coagulant, and snap frozen. Lymph nodes and spleens were processed into single cell suspensions, then analyzed by flow cytometry. Lungs were collected and weighed prior to fixation in formalin for 24 hours then transferred to 70% ethanol. Tumors were collected and fixed in formalin/70% ethanol for analysis.

### Pharmacodynamic studies

The pharmacodynamic effects of the test compounds SB 11285, and SB 11312 (3 mg/kg, I.V., on day 1 and day 5) were conducted in BALB/c mice bearing an average of 100 mm^3^ CT26 tumors. The animals were randomized into multiple groups based on tumor volume. The anti-tumor activity was evaluated as a measure of the decrease in tumor volume in treated animals compared to the control animals. On day 6, post-dose, the study was terminated, and animals were humanely euthanized. Gross pathological observation was carried out to evaluate any abnormality. Blood, lymph nodes, spleen and tumor were collected, and flow cytometric analysis was carried out. Plasma was used for multiplex analysis for cytokine release whereas tumors were also collected and fixed in 10% formaldehyde for IHC analysis.

### Immunophenotyping

Isolated single cell suspensions from tumor, spleen, and lymph nodes were seeded (0.4 x 10^6^ cells per 50 μL per well) onto 96 well U bottom plates. 50 μL of the required antibodies (2 X concentration) were added to the appropriate wells, mixed, and incubated for 30-45 minutes in ice. Cells were washed in Cytofix/perm buffer once and re-suspended in Cytofix/perm buffer for 20 minutes at RT. Cells were washed once in 1x perm buffer and re-suspended in perm buffer for 20 minutes. Cells were incubated with FoxP3 antibody for 30 minutes and washed. Cells were re-suspended in staining buffer and acquired in BD FACSVerse flow cytometer with appropriate compensation controls. Immunophenotyping was performed using the markers given below:

- T cells: CD3+CD4+ or CD3+CD8+
- Treg: CD4+ CD25+ Foxp3+
- Granulocytic MDSC: CD11b+ Ly6G
- Monocytic MDSC: CD11b+ Ly6C

### Immuno-histochemistry

Paraffin-embedded tissue sections were de-waxed in xylene, washed in a graded alcohol system, and the peroxidase activity was quenched with 3% hydrogen peroxide for 10 minutes to avoid non-specific staining. The sections were washed three times with PBS followed by incubation overnight with anti-CD8 antibody and anti-F4/80 antibody (Abcam) at 1 in 100 dilutions at 4°C. The sections were then washed with PBS and incubated at room temperature for approximately 20 minutes with a reaction enhancer kit followed by three washes in PBS, incubation with secondary antibody at room temperature for 20 minutes and staining with 3,3-diaminobenzidine. The sections were dehydrated and sealed after redyeing with hematoxylin.

## Results

### Library Synthesis and Discovery of Leads

Our strategy for the discovery of STING agonists involved screening a library of Pu-Py CDNs (Fig. 1) in cell-based assays to identify compounds that induced Type I interferons (IFN) in human cells through the activation of the STING signaling pathway. We synthesized a library of over 100 cyclic dinucleotides by varying purine-pyrimidine nucleobase combinations or substitutions in the ribose rings and incorporating mixed phosphodiester-phosphorothioate (PS) internucleotidic connectivities, in which the single PS-linkage was selectively S-alkylated using a multitude of alkyl, aryl, and ar-alkyl substituents (42).

**Fig.1.**
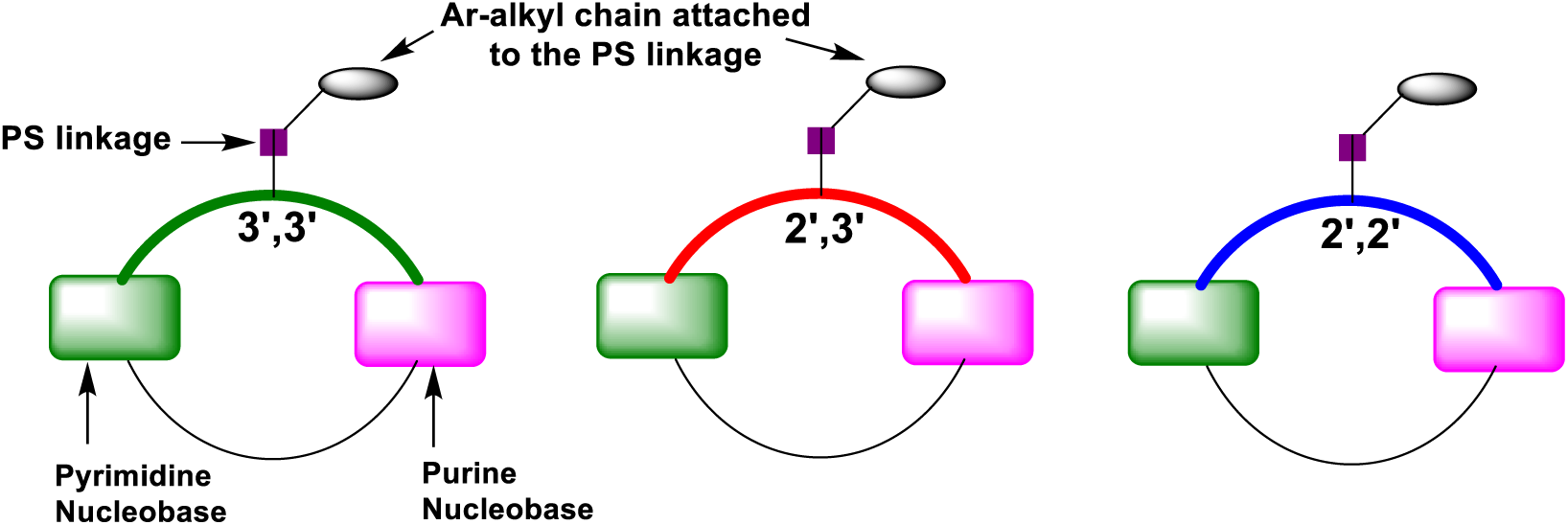
General structural representation of diverse Purine-Pyrimidine classes of 3’5’-cyclic dinucleotides (CDNs). Note that the 3’,3’ class of CDNs is also referred to as 3’,5’-CDNs which signifies that the internucleotidic connectivities are from 3’-hydroxyl to 5’-hydroxyl through phosphodiester (PO) or phosphorothioate (PS) linkages. In each case, the ar-alkyl chain is attached to the PS backbone of the CDN through chemo selective *S-*alkylation.

Using the HEK293-based IFN-responsive stable luciferase cell line, called SZ14, we identified and selected initial leads as “hits” that showed > 5-fold induction of IRF3-ISG54 compared to untreated cells. These early lead compounds were then used as scaffolds for the design and preparation of a focused library of compounds for further screening and lead optimization.

### EC_50_ assessment and Lead Optimization for the Discovery of SB 11285

Following the identification of hits and the synthesis of analogs, we proceeded with the utilization of SZ-14 reporter cells to detect potent compounds. Next, promising lead compounds were further tested as STING agonists in THP1-Dual-WT cells and THP1-Dual-STING KO cells (InvivoGen). These cells belong to the human monocytic cell lineage and express two inducible reporter genes, facilitating concurrent assessment of the IRF and NF-κB pathways **(Fig. S1)**. Through an iterative process of synthesis and screening of several hundred compounds of the 3’,5’-PuPy-CDN family, we identified SB 11285 as the lead compound. **Table 1** shows the EC_50_ values of SB 11285 for IRF and NF-κB induction. SB 11285 is a 1:1 diastereomeric mixture of two isomers, SB 11326, and SB 11327, both of which exhibited similar STING agonist activities in the assays. Additionally, SB 11312, the Pu-Py CDN without the ar-alkyl substitution and the major metabolite of SB 11285, was found to be a STING agonist, but with a decreased EC_50_ of 3 µM compared to SB 11285. In these assays, 2’,3’-cGAMP, the natural STING agonist, showed EC_50_ for IRF induction of 1.3 µM without any NF-κB activity.

**Table 1.**
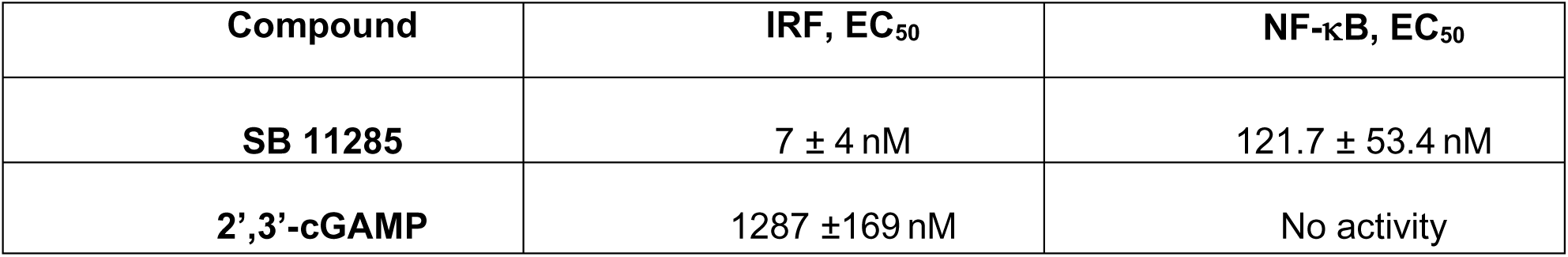
EC_50_ values of SB 11285 and cGAMP as % IRF and NF-κB induction in THP-1 cells.

SB 11285 and its analogs were evaluated in cell-based assays without the aid of transfection reagents, while cGAMP was tested using transfection reagents such as lipofectamine or digitonin. Under these assay conditions, SB 11285 exhibited superior activation of STING in various human immune cells, including PBMCs, monocytes, and monocyte-derived dendritic cells, as well as, in mouse splenocytes and bone marrow-derived macrophages, compared to other STING agonists such as cGAMP and ADU-S100 (Table 5, Table S4).

Utilizing flow cytometric analysis, it was observed that the biotin-tagged SB 11285 compound was preferentially taken up by myeloid dendritic cells and monocytes within PBMCs, while showing minimal uptake by B cells or T cells (**Fig**. **S2**). The selective uptake of SB 11285 might involve mechanisms such as phagocytosis or micropinocytosis (43) and require further investigation.

### Estimation of KD of SB 11285 by Surface Plasmon Resonance (SPR)

A biotin-tagged SB 11285 was used in a SPR protocol to demonstrate its binding to WT-hSTING-CTT. Using this approach, the equilibrium dissociation constant (KD) of binding of SB 11285 to hSTING-CTT was estimated to be ca.122 nM (**Fig. S3**).

### SB 11285 activates the STING Signaling Pathway

As mentioned earlier, the binding of CDNs to STING triggers the activation of the STING-TBK1 complex, leading to the translocation of STING from the ER to perinuclear endosomes. This process additionally involves the palmitoylation of STING and subsequent TBK1 phosphorylation. Activated STING then induces nuclear translocation of IRF3 and initiation of transcription resulting in IFNβ expression, NF-κB activation, and gene expression of cytokines (12). SB 11285 demonstrated potent induction of IRF3 nuclear translocation in THP1-derived macrophages (**Figs. S4, S5**), indicating the activation of the type I IFN signaling pathway.

STING activation relies on its palmitoylation at Golgi (Cys88/91), enabling downstream signaling and IFN expression (46). The palmitoylation inhibitor 2-bromo-palmitic acid (2-BP) blocked the SB 11285-induced type I IFN response and IFN-γ production from NK cells (data not shown).

To identify intermediary proteins in the IFN signaling cascade following the activation of STING, Western blot analysis of THP-1 cells treated with SB 11285 was undertaken which revealed that the phosphorylation of TBK1 and IRF3 occurred within 2 and 4 hours, respectively, followed by pIκBα formation within 8 hours indicative of NF-κB activation. Additionally, SB 11285 induced STING degradation within 24 hours, potentially through negative feedback of the STING pathway following activation **(Fig S5)**.

The STING agonist activity of SB 11285 was also assessed using human THP-1 dual parental cell lines and human monocytic THP-1 cells with an IRF-inducible luciferase reporter, both expressing STING-HAQ, alongside the corresponding STING-KO line. Cells were treated with SB 11285 and cGAMP in a 3-fold dilution, 9-point dose-response from 0.0076 µM to 50 µM for 24 hours (in triplicate). SB 11285 selectively activated the IRF-inducible luciferase reporter in THP-1 Dual parental and STING-HAQ engineered lines with a normalized EC_50_ of 0.085 µM, showed complete loss of activity in the STING-KO line. This confirmed that SB 11285 induced IRF signaling in a STING-dependent manner with superior potency compared to cGAMP and ADU-S100 (**Table 2)**.

**Table 2.** Activation of IRF3 by SB 11285 in human THP-1 cells is STING-dependent.

| Compound | Human THP-1 cell line – IRF3 reporter activation |  |  |  |
| --- | --- | --- | --- | --- |
|  | STING-HAQ |  | KO-STING |  |
| | $\text{EC}_{50}$ ($\mu\text{M}$) | Maximum fold-change | $\text{EC}_{50}$ ($\mu\text{M}$) | Maximum fold-change |
| **SB 11285** | $0.085 \pm 0.003$ | $432 \pm 8$ | $>10$ | $1 \pm 0$ |
| **cGAMP** | $>10$ | $137 \pm 5$ | $>10$ | $1 \pm 0$ |
| **ADU-S100** | $3.0 \pm 0.4$ | $436 \pm 13$ | $>10$ | $1 \pm 0$ |

STING signaling by SB 11285 was further assessed in RAW Macrophages using RAW-Lucia ISG cells expressing mSTING-WT, mSTING-KO, TBK1-KO, IRF3-KO, and IRF7-KO. Following the treatment of cells with SB 11285 or DMSO, IRF induction was either abolished or significantly reduced in mSTING-KO, TBK1-KO, IRF3-KO, and IRF7-KO cells compared to SB 11285-treated STING-WT cells **(Fig. S6)**. This underscores the specificity of SB 11285 for STING and highlights the involvement of intermediary proteins in the STING signaling cascade.

### SB 11285 Causes Apoptosis by Modulating Pro-, and Anti-apoptotic Molecules

STING activation has been linked to apoptosis induction in cancer cells via apoptosis regulation pathways including Endoplasmic Reticulum Stress, NOD-like receptor and NF-κB pathways, as well as, IRF3-BAX interactions (47–50). Despite the reported downregulation of hSTING in various cancers (48), we detected its expression in human lung, prostate, ovarian, and colon cancer cells. Therefore, we investigated whether SB 11285-induced hSTING activation could result in apoptosis in these cancer cells. Gene expression analysis revealed a dose-dependent increase in apoptosis in mouse A20 lymphoma cells treated with SB 11285, accompanied by an elevated oncogene-derived proteins Bax/Bcl2 ratio indicative of apoptosis promotion **(Figs. S7, S8)**. These findings underscore highly selective STING agonism by SB 11285 in inducing IFN expression in human and mouse cells while triggering apoptosis in tumor cells in a STING-dependent manner (51–55).

### SB 11285 Activates Multiple hSTING Alleles and is More Potent Than cGAMP

Polymorphisms in the hSTING gene affect responsiveness to bacterial-derived CDNs (19, 27). Five haplotypes (WT, REF, HAQ, AQ, and Q alleles) vary in amino acids at positions 71, 230, 232, and 293 (54,55). SB 11285 demonstrated potent activation of all hSTING alleles compared to cGAMP, albeit with diminished activity in R232H and R232Q alleles compared to WT-hSTING (**Fig. S9**).

### SB 11285 Causes the Induction of Pattern Recognition Receptors

Innate sensing of foreign nucleic acids triggers immune responses and RIG-I and STING pathways are involved in RNA and DNA sensing, respectively, with significant crosstalk between various PRRs (56). SB 11285, a STING activator, induced RIG-I pathway genes in THP-1 cells but not in PBMCs, unlike cGAMP which induced PD-1 and PD-L1 (**Fig. S10**). The significance of the selective induction of PRRs by SB 11285 in these cells is not known and requires further investigation.

### SB 11285, a Canonical 3’,5’-Pu-Py CDN, is More Potent Than Non-canonical CDNs

We compared the activation of IRF3 and NF-κB pathways by SB 11285 with other STING ligands in THP-1 cells expressing hSTING-HAQ and wild-type hSTING (WT-hSTING) (**Table 3, 4**). SB 11285 showed potent activation of both pathways in both cell lines, significantly outperforming cGAMP and ADU-S100 in potency for both pathways and in both cell lines. SB 11285’s potency was markedly higher, with an EC_50_ for STING-HAQ (>118-fold and 35-fold lower than cGAMP and ADU-S100, respectively). Moreover, SB 11285’s agonist activity was dependent on STING, as evidenced by the abolishment of IRF and NF-κB induction in KO-STING cells **(Table 2)**.

**Table 3.** SB 11285 activates the IRF3 and NF-κB pathways in human STING-HAQ THP-1 cells.

| Compound | Human THP-1 cells (STING-HAQ) |  |  |  |
| --- | --- | --- | --- | --- |
|  | IRF3-reporter |  | NF-κB-reporter |  |
|  | EC<sub>50</sub> (μM) | Maximum fold-change | EC<sub>50</sub> (μM) | Maximum fold change |
| **SB 11285** | 0.13 ± 0.01 | 278 ± 69 | 0.37 ± 0.03 | 10 ± 1 |
| **cGAMP** | >10 | 53 ± 26 | >10 | 1 ± 0 |
| **ADU-S100** | 8.9 ± 0.1 | 202 ± 63 | >10 | 2 ± 1 |

**Table 4.** SB 11285 activates the IRF3 and NF-κB pathways in human STING-WT THP-1 cells. **Please note.** Data in Tables 1-4 are presented as mean values ± SD from minimum three replicates and three independent experiments. Fold-change is expressed relative to the no compound control.

| Compound | Human THP-1 cells (STING-WT) |  |  |  |
| --- | --- | --- | --- | --- |
|  | IRF3-reporter |  | NF-κB-reporter |  |
|  | EC<sub>50</sub> (μM) | Maximum fold-change | EC<sub>50</sub> (μM) | Maximum fold change |
| **SB 11285** | 0.5 ± 0.1 | 179 ± 21 | 0.66 ± 0.04 | 6 ± 1 |
| **cGAMP** | >10 | 20 ± 3 | >10 | 1 ± 0 |
| **ADU-S100** | >10 | 52 ± 7 | >10 | 1 ± 0 |

In summary, in hSTING-WT cells, EC_50_ values of SB 11285 for IRF3 pathway activation was >20-fold lower than cGAMP and ADU-S100 while for NF-κB pathway activation was >14-fold lower than cGAMP and ADU-S100. In hSTING-HAQ cells, the EC_50_ value of SB 11285 for IRF3 pathway activation was ≥ 89-fold lower than cGAMP and ADU-S100. The EC_50_ value of SB 11285 for NF-κB pathway activation was >25-fold lower than cGAMP and ADU-S100.

### Molecular Modeling and Dynamics Simulation of Binding of SB 11285 with STING Identify Critical Components for Interaction

To understand the molecular interactions and the binding modes of the 3’,5’-PuPy-CDNs, we performed computational modeling (Schrodinger Suite) and molecular dynamics (MD) simulation (Amber18 package) (350 ns) of SB 11285 using published crystal structures of cGAMP bound to hSTING [PDB ID: 4F5D] (24–27) and found that while key residues Tyr167 and Arg238 within CDN pocket helped the binding of SB 11285 to hSTING, additional interactions between the aryl-alkyl chain of SB 11285 with the amino acid residues from the gatekeeper loop of B-chain dimer enhanced the further stabilization of STING dimer interface and tighter binding based on MD simulations and earlier reported simulation of C-terminal tail [CTT] structure of hSTING (60). It is expected that the terminal end of Ar-alkyl chain of SB 11285 may additionally interact with the C-terminal tail (CTT) of hSTING.

Attempts to obtain the crystal structure of SB 11285 with hSTING-CTT have so far been unsuccessful.

Based on molecular modeling and MD, we generated several hSTING mutants and evaluated them in cell-based assays using plasmids transfected into HEK293T cells. Specifically, mutation of either Tyr167 or Arg238 residues abolished hSTING activation by SB 11285 (**Fig. S11**), indicating the involvement of nucleobases of SB 11285 in Tyr167 or Arg238 binding, and recognition by hSTING, which is similar to that observed with cGAMP interactions with hSTING (26, 27).

### Selectivity Assays Demonstrate that SB 11285 is a Highly Specific STING Agonist

SB 11285’s selectivity and specificity for STING were confirmed by evaluating its activity in THP-1 cell lines lacking cGAS or IFI16 expression, the two pattern recognition receptors that activate the STING pathway. SB 11285 showed comparable activity in cells with or without cGAS and IFI16 expression, indicating that it does not activate these receptors or pathways (**Table S1**).

Additionally, SB 11285 did not stimulate any of the tested toll-like receptors (TLRs), NOD-like receptors (NLRs), C-type lectin receptors (CLRs), or RIG-I like receptors (RLRs) in HEK293 cell lines expressing these receptors, further confirming its specificity for STING-induced activation of the IFN signaling cascade (**Table S2**).

### SB 11285 Activates Mouse STING More Potently Than Non-canonical STING Ligands

SB 11285 activated mouse STING (mSTING) in murine B16 melanoma cells, as evidenced by its ability to induce IRF3-reporter gene activation in wild-type B16 cells but not in KO-mSTING cells (**Table S3**). SB 11285 exhibited higher potency in activating the IRF3 pathway compared to cGAMP and ADU-S100 in wild-type B16 cells. These results supported the rationale for in vivo studies of SB 11285 in syngeneic mouse tumor models. Additionally, SB 11285 demonstrated the activation of Type I, Type II, and Type III IFN responses, surpassing other STING ligands in potency (**Fig. S12**).

### Comparative Profiling of SB 11285 and Non-canonical STING Ligands in Human and Mouse Primary Immune Cells

SB 11285 induced a dose-dependent production of various cytokines in both human PBMCs and mouse splenocytes, including type I IFNs, pro-inflammatory cytokines, anti-inflammatory cytokines, and chemokines. The potency varied across different cytokines and cell types, with SB 11285 demonstrating comparable or higher potency compared to cGAMP and ADU-S100 in inducing cytokine production. Overall, SB 11285 activated both the IRF3 and NF-κB pathways in a STING-dependent manner in human PBMCs and mouse splenocytes (**Table 5, Table S4**).

**Table 5.** Summary of cytokines induced by SB 11285 in healthy human donor PBMCs.

| Cytokine |  | SB 11285 |  | cGAMP |  | ADU-S100 |  |
| --- | --- | --- | --- | --- | --- | --- | --- |
| | | EC<sub>50</sub>(μM) | Max fold-change | EC<sub>50</sub>(μM) | Max fold-change | EC<sub>50</sub>(μM) | Max fold-change |
| Type I IFN | IFN-α | 0.9 ± 0.7 | 560 ± 575 | >10 | 31 ± 24 | 2.6 ± 2.9 | 582 ± 242 |
| | IFN-β | 0.3 ± 0.2 | 36 ± 23 | >10 | 1 ± 0 | >10 | 14 ± 8 |
| Type II IFN | IFN-γ | 1.0 ± 0.4 | 836 ± 891 | >10 | 9 ± 3 | >10 | 392 ± 374 |
| Immuno-modulatory | IL-12p70 | 0.8 ± 0.3 | 44 ± 64 | >10 | 1 ± 0 | >10 | 7 ± 2 |
| | IL-18 | 0.7 ± 0.1 | 15 ± 10 | >10 | 1 ± 0 | >10 | 8 ± 7 |
| | GM-CSF | 0.4 ± 0.3 | 12 ± 3 | >10 | 2 ± 1 | 2.9 ± 0.4 | 21 ± 2 |
| Pro-/Anti-inflammatory | IL-1α | 0.4 ± 0.2 | 38 ± 37 | >10 | 13 ± 11 | 2.6 ± 0.2 | 52 ± 8 |
| | IL-1β | 1.0 ± 0.8 | 486 ± 456 | >10 | 21 ± 11 | >10 | 323 ± 309 |
| | IL-6 | 0.3 ± 0.2 | 155 ± 137 | >10 | 30 ± 20 | 2.3 ± 0.6 | 213 ± 183 |
| | TNF-α | 0.9 ± 0.5 | 564 ± 302 | >10 | 4 ± 3 | >10 | 174 ± 141 |
| | IL-10 | 0.03 ± 0.03 | 8 ± 2 | >10 | 4 ± 3 | 0.8 ± 0.6 | 7 ± 3 |
| **Chemokines** | **IP-10** | 0.03 ± 0.01 | 10 ± 7 | 1.8 ± 1.0 | 4 ± 2 | 0.4 ± 0.3 | 8 ± 5 |
|  | **MIP-1α** | 0.05 ± 0.04 | 204 ± 179 | >10 | 51 ± 38 | 1.1 ± 0.6 | 273 ± 234 |
|  | **MIP-1β** | 0.06 ± 0.09 | 91 ± 69 | >10 | 72 ± 77 | 1.2 ± 0.8 | 118 ± 82 |
|  | **RANTES** | 0.08 ± 0.06 | 19 ± 19 | 1.6 ± 1.4 | 5 ± 3 | 0.7 ± 0.5 | 6 ± 3 |

Human PBMCs from 4 to 6 healthy volunteers were stimulated with SB 11285, cGAMP and ADU-S100. Cytokine responses were measured after 24 hours using a 45-plex Luminex assay and ELISA for IFN-β. Mean values ± SD from 4 to 6 n = 3 to 6 independent donors. Fold-change is expressed relative to the no compound control.

### Pharmacodynamics of SB 11285 in Tumor-bearing Mice

We ascertained the effect of SB 11285 and its metabolite SB 11312 on the induction of cytokines in BALB/c mice bearing CT26 syngeneic tumors. Treatment with SB 11285, or SB 11312 (3 mg/kg, I.V., on day 1 and day 5) resulted in significant inhibition of tumor growth by day 6 with TGI being ca. 92% for SB 11285 and 67% for SB 11312 as compared to the vehicle control animals (p<0.001).

There was a statistically significant elevation in IFNα/IFNβ cytokines in SB 11285-treated animals on day 1, 4hrs post dose (p<0.001), as well as, significant elevation (p<0.0001) of TNF-α, IL-6, KC, MCP-1, IL12p70, MIP-2 and IL-10 in blood from animals treated with SB 11285, but not SB 11312, as compared to vehicle control group on day 1 (**Fig. 2**). On day 6 (24 hrs., after dosing), SB 11285-treated animals showed a statistically significant elevation in TNF-α, MCP-1, IL-10, MIP-2, and IL-6 in blood compared to untreated animals, however, the levels of the cytokines were significantly lower when tested 24 hours post dosing when compared t4 hours post dose testing suggesting that the cytokine response peaks within a few hours after dosing of SB 11285 and SB 11312. No elevation in IFNα was observed on day 6 with SB 11285 treatment. A small increase in IFNβ induction in response to SB 11285 and SB 11312 was observed on day 6. These results could be because the induced levels may not be enough to stimulate a positive feedback loop mediated by the IFNβ-IRF7-IFNα/β pathway and may explain why IFN-α levels are not induced by SB 11285 or SB 11312 on day 6. It is also possible that the IFN response could peak within a few hours after dosing of SB 11285 and SB 11312 and may not be detected 24 hours after dosing. SB 11312-treated animals showed a statistically significant increase in IFNs, KC, MIP-2, TNFα, and IL-10 on day 6 (**Fig. 2**).

**Fig. 2.**
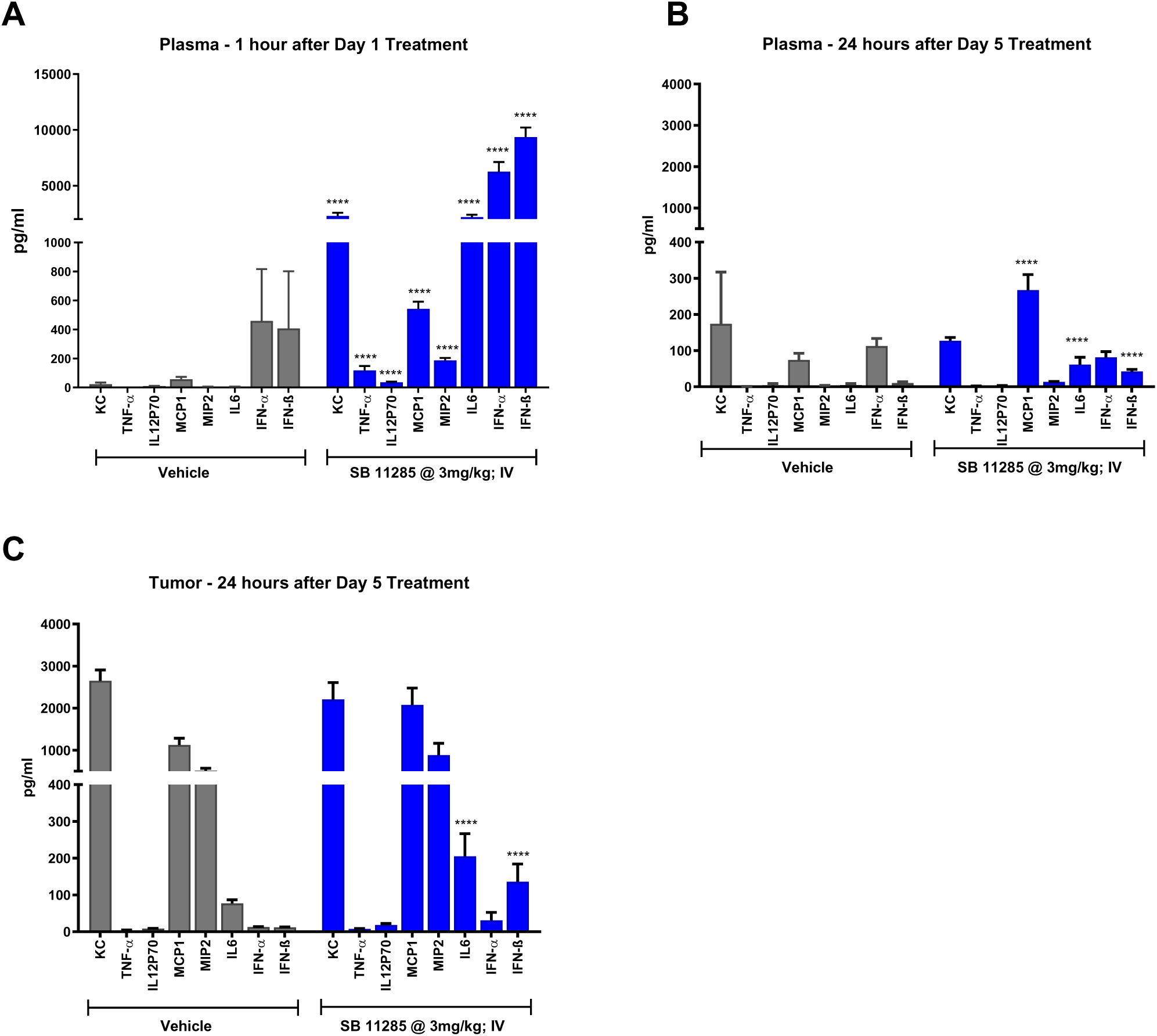
Pharmacodynamic effects of SB 11285 & SB 11312 in BALB/c mice bearing subcutaneous syngeneic CT26 tumor. The pharmacodynamic effect of SB 11285 and its metabolite SB 11312 were conducted in BALB/c mice bearing CT26 syngeneic tumor. The animals were randomized into 6 treatment groups (3 mg/kg, I.V., on day 1 and day 5). On day 1, 1.5 hr., post-treatment and on day 6 (24 hrs., post-treatment on day 5), blood samples were collected from all animals and plasma was separated for further analysis. On day 6, the study was terminated, and blood, lymph nodes, spleen and tumor were collected, and flow cytometric analysis was carried out. **(A).** Plasma was used for multiplex analysis by performing the mouse magnetic Luminex assay. The plasma levels of IFNα and IFNβ were quantified using the mouse IFNα/β procarta plex. **(B). The assessment of IFNs, cytokines, chemokines, and T cells in tumor site**. Tumor tissues were cut into small pieces and dissociated using gentleMACS dissociator, incubated with enzyme mix for 40 minutes at 37 °C and centrifuged. Single cells isolated from tumor tissues were pelleted, homogenized, and supernatants were analyzed for cytokines using Luminex mouse magnetic assay and mouse IFNα/β procarta plex. Total protein estimation was performed in the tumor homogenates using Pierce BCA protein assay reagent. Flow cytometric analysis was done to quantify Tregs in tumor and lymph nodes.

Treatment with SB 11285 also showed a significant increase in the levels of IFN-α and IFN-β in the tumor homogenates compared to vehicle treated group (p = <0.05). In the case of tumor cells, high levels of KC and MCP-1 levels were observed in the vehicle-treated group, whereas the treatment with SB 11285 or SB 11312 showed moderate increase in cytokines TNFα, IL-6, MCP-1, IL-1β, MIP-2, IL12p70, IL-10 and IFNα (**Fig. 2**), but was not statistically significant. It is to be noted that the cytokines measured in the tumor homogenates may be produced by the tumor cells themselves or other components of the TME including the immune cells and endothelial cells. Future analysis of immune cells isolated from the tumors may provide more information about the cytokine induction/activation of specific tumor-infiltrating immune cells in response to SB 11285 and SB 11312 treatment.

Treatment with SB 11285 resulted in a significant increase (p < 0.01) in CD3+CD45+ve T cells in the peripheral blood, a decrease in CD3+CD45+ve T cells in tumor (p < 0.001) and a significant increase in CD45+ T cells in the lymph nodes. Treatment with SB 11312 also resulted in a highly significant increase in CD3+CD45+ve T cells in blood (p < 0.0001), and a significant decrease of CD45+ve cells in the spleen (p < 0.01). SB 11312 did not induce T cell infiltration into the tumor.

Since both SB 11285 and SB 11312 caused significant tumor regression by day 6 in the CT26 model, the lack of CD8 +T cells in tumors may reflect that in these studies, the timing of collection of peripheral cells, and tissues may need further optimization to understand the kinetics of activation and migration of T cells from periphery to the tumor site. It is also likely that once the tumor regresses significantly, the CD8 +T cells may not persist at the tumor site but instead may circulate back into the periphery as part of natural defense mechanism involving tumor surveillance.

Tumor-infiltrating Tregs play direct roles in promoting immune evasion, and the development of a pro-tumorigenic TME (9). In this study, treatment with SB 11285 significantly reduced the Treg infiltration in the tumor (p < 0.001) further suggesting the role of SB 11285 in promoting immune activation and enabling anti-tumor immunity. Interestingly, an increased number of Treg cells were observed in the periphery following SB 11285 treatment. Treatment with SB 11312 also showed moderate reduction of Tregs in tumor (**Fig. 2**).

MDSCs are a heterogeneous population of immature myeloid cells that include monocytic (mMDSC) and granulocytic (gMDSC) subsets both of which have been shown to be immuno-suppressive. MDSCs and Tregs can co-operatively function to promote tumor-immune evasion (8, 9). Indeed, treatment with SB 11285 showed significant reduction in granulocytic (CD11b+Ly6G+) and monocytic (CD11b+Ly6C+) MDSCs in peripheral blood. In the tumor, SB 11285 significantly elevated the number of granulocytic MDSCs and reduced the number of monocytic MDSCs. Treatment with SB 11312 did not show any statistically significant changes of these cells in the tumor, but significant reduction was observed in granulocytic MDSCs in peripheral blood. The tumor regression seen with SB 11285 and the reduction in the monocytic MDSCs may suggest that these cells may be the primary immunosuppressive MDSCs or M2 macrophages.

### Antitumor effects of SB 11285 are mediated by CD8+ T cells

To further assess the role of T-cells and evaluate whether treatment with SB 11285 could induce an enhanced tumor-specific IFNγ T cell response for anti-tumor activity in the CT26 colon carcinoma mouse model, spleens were harvested after treatment of a single dose of SB 11285 or cGAMP and were tested in IFNγ ELISPOT assay with irradiated tumor cells. At 2.6 mg/kg I.T., dose, SB 11285 was comparable to cGAMP in inducing IFNγ-secreting T cells (**Table 6**) with the average IFNγ spots per 10^6^ splenocytes ± SEM was 581±39, and 625±59 for SB 11285, and cGAMP, respectively. At 5.3 & 10.6 mg/kg dose SB 11285 I.T., the average IFNγ spots per 10^6^ splenocytes ± SD were 715±17 and 589±80, respectively. In the case of single dose I.V., with SB 11285, administered at 1.1, 3.2 & 5.3 mg/kg in the CT26 model significant presence of IFNγ spots per 10^6^ splenocytes was observed, respectively (**Table 6**) showing robust induction of IFNγ in splenocytes.

**Table 6.**
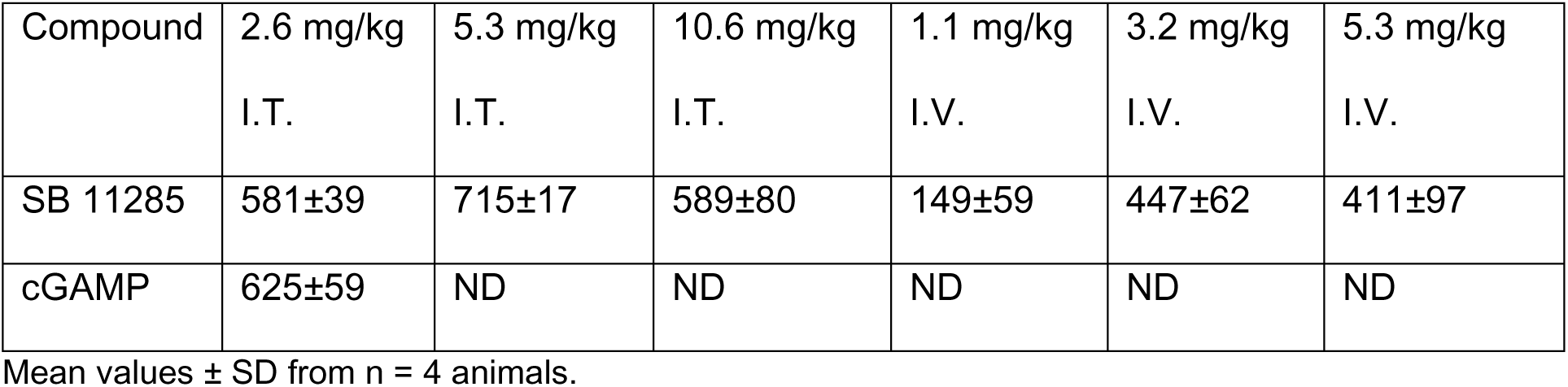
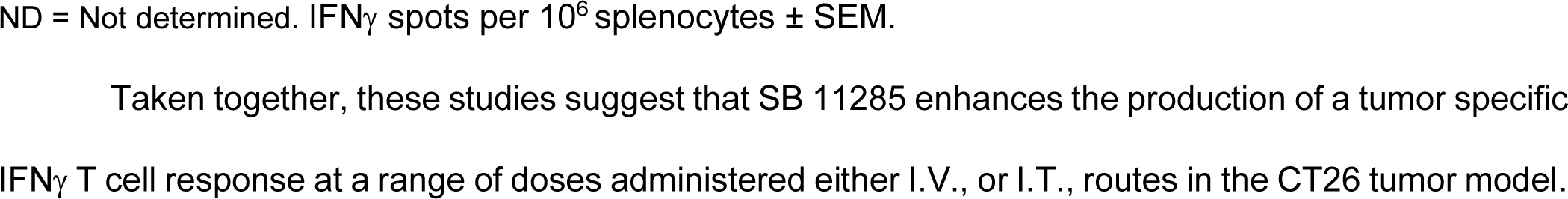
Tumor-specific IFNγ T cell response in CT26 after SB-11285 treatment.

Finally, to further elucidate the role of CD8+ T cells in the antitumor immunity provided by I.V. administered SB 11285, we evaluated the anti-tumor activity of SB 11285 alone or in combination with anti-CD8b antibody (that causes depletion of CD8 T cells) in BALB/c mice bearing subcutaneous syngeneic CT26 tumors. Whereas I.V. administration of SB 11285 resulted in significant anti-tumor activity (TGI, 96%), combination of SB 11285 with anti-CD8b antibody resulted in significant reduction of antitumor activity (TGI 40%) **(Fig. 3)**. These data strongly suggest that CD8 T cells play an important role in vivo in SB 11285-mediated antitumor activity.

**Fig. 3.**
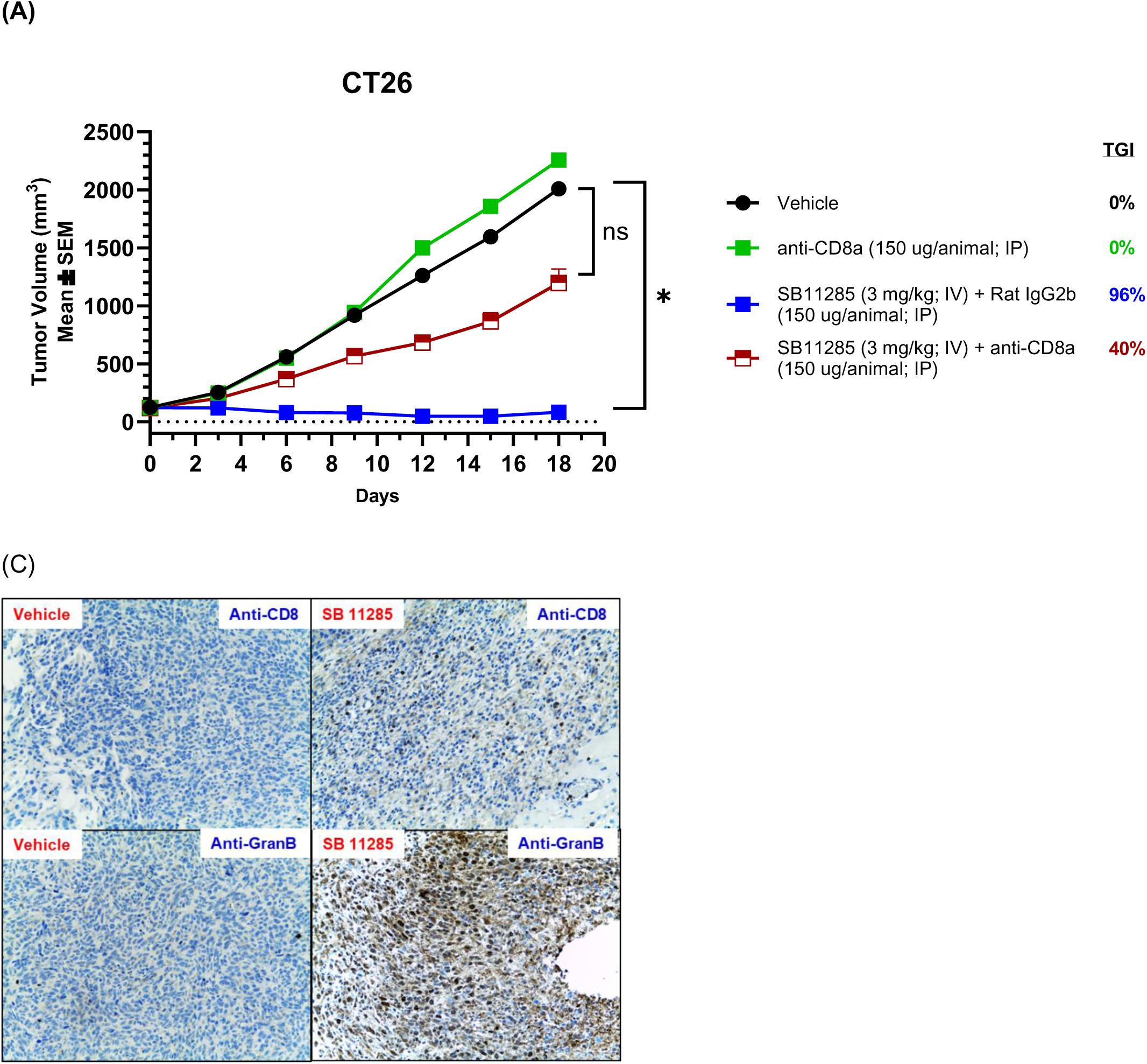
Evaluation of the anti-tumor activity of I.V., SB 11285 + isotype control alone and in combination with anti-CD8b in BALB/c mice bearing subcutaneous syngeneic CT26 tumors. **(A)** Groups of 8 mice were subcutaneously administered 2 X 10^6^ CT-26 murine colorectal carcinoma cells and tumors were allowed to grow until MTV was approximately 100 mm^3^. SB 11285 was administered on days 1, 5, 9, and 14 alone, or in combination with anti-CD8b antibody and rat IgG2b antibody on days −1, 0, 2, 6, 10, 14 and 18. MTVs were measured on indicated days and plotted as shown. **(B)** Mouse tumor volumes plotted as mean ± SD, on day 14 following treatment with Rat IgG2b, or SB 11285 + Rat IgG2b or SB 11285 + Anti-CD8b. Statistical significance was assessed by unpaired t-test, two tailed. (**C**) Tumor tissue was collected on day 6 to analyse Cytotoxic lymphocyte cell population by using anti-granzyme antibody staining in IHC samples.

### I.T., SB 11285 alone, and in combination with cyclophosphamide Induce Profound Anti-Tumor Effects and Immune Memory in Mouse Tumor Models

To evaluate whether the combination with anticancer drugs can potentiate or inhibit the anti-tumor activity of SB 11285, we carried out studies with cytotoxic drugs such as cyclophosphamide (CP). The efficacy studies of SB 11285 alone, and in combination with the cytotoxic with cyclophosphamide were conducted in the A20 lymphoma syngeneic mouse model. Thus, following the establishment of the subcutaneous tumors with an average size of 100mm^3^ in mice, I.T., administration of SB 11285 alone at a dose of 5 mg/kg, and in combination with CP at 100 mg/kg were initiated. By day 15, highly significant TGI of 86%, 100%, and 93% were seen in the SB 11285, cyclophosphamide (CP), and the combination treatment groups respectively, compared to the vehicle control groups (*P* < 0.001), and all animals were followed for TGD phase for further 72 days (**Table 7**, **Fig. 4**) which gave TGD of 64, 156 and 288% respectively. The enhanced antitumor activity of SB 11285 when combined with the cytotoxic drug, cyclophosphamide, illustrates the benefit of the combination therapy of SB 11285 with cytotoxic agents.

**Fig. 4.**
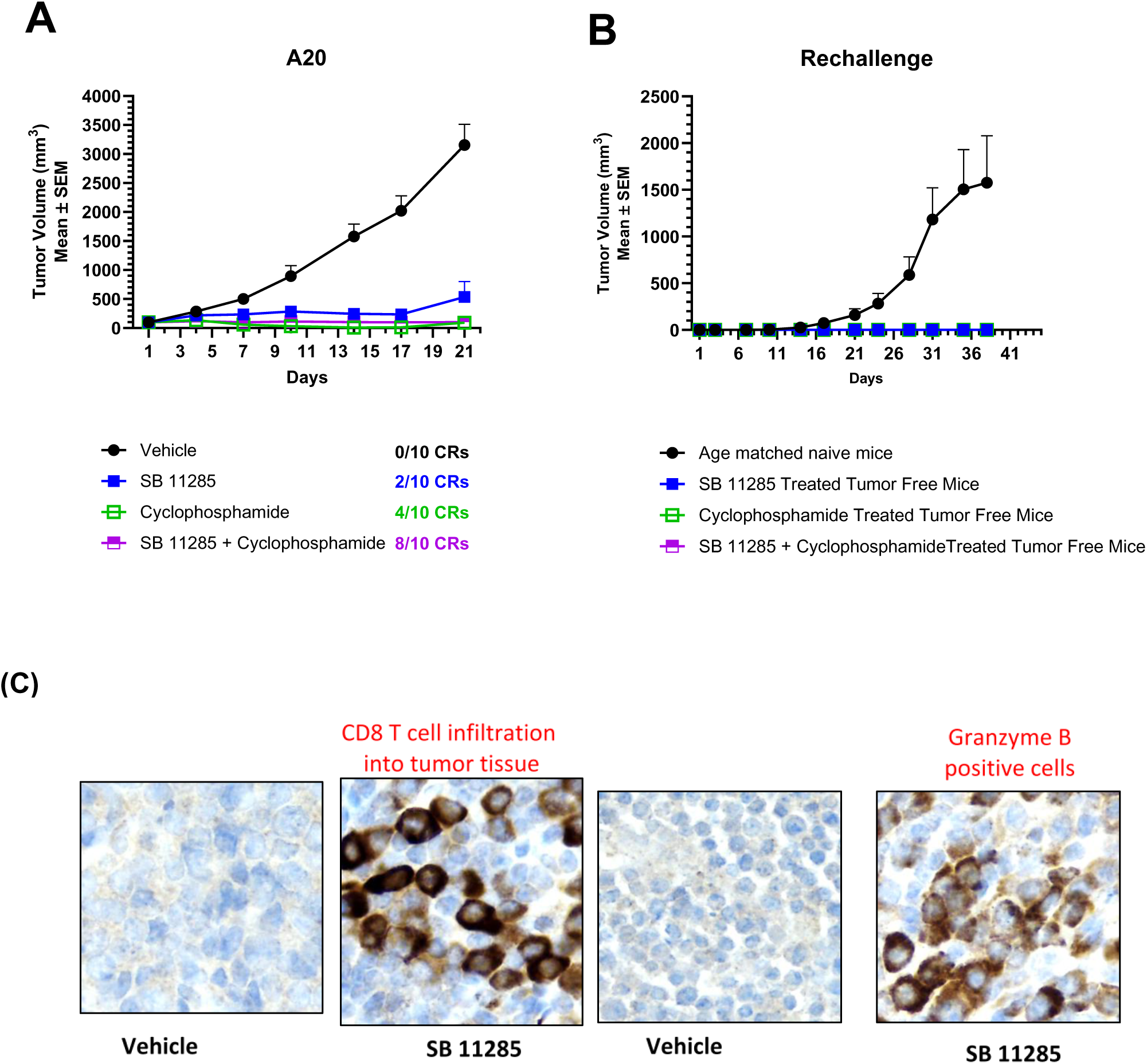
Evaluation of SB 11285 in Syngeneic A20 Lymphoma Model: Briefly, A20 cells were implanted subcutaneously in the right flank region of female BALB/c mice (n = 10). Dosing was initiated when the average MTV reached 100 mm3. SB 11285 was administered intratumorally (I.T.) on days 3, 4, 6, 8 and 10 at 100 µg in 50 µl in saline. Cyclophosphamide (CP) was administered intraperitoneally (I.P.) two times on days 1 and 2 at a dose of 100 mg/kg. (A). Tumor volumes measured on day 22 after the initiation of treatment. Response summary of the TGI and TGD data in A20 lymphoma model is shown in Table 7. (C). Detection of CD8+ T cells and NK cells in tumor tissues. Immunohistochemistry of tumor tissues was carried out using anti-CD8 Antibody and anti-Granzyme B antibody one day after end of dosing (day 11) to evaluate infiltration of CD8 T cells and cytotoxic cells respectively into tumor tissue.

**Table 7.** Summary results following different treatments in the syngeneic mouse A20 tumor model.

| Group | MTV<br>(day36) | MTV<br>(day71) | Partial<br>Responders | Complete<br>Responders | %<br>TGD |
| --- | --- | --- | --- | --- | --- |
| Vehicle | 3155 |  | 0 | 0 | 0 |
| SB 11285 | 534 | 0 (4) | 1 | 3 | 64 |
| CP | 92.4 | 0 (4) | 5 | 4 | 156 |
| CP+11285 | 101.5 | 0 (9) | 1 | 9 | 288 |

In this study, tumor tissues were also analyzed for the presence of CD8+ T cells, macrophages, and NK cells. Thus, tumor cells were collected one day after the end of the treatment (on day 11) in the SB 11285 alone treatment group, and the paraffin sections were stained with anti-CD8, and anti-Granzyme B antibodies, which revealed the presence of significant number of CD8+ T cells, and NK cells in the TME of SB 11285-treated animals compared to that of the untreated controls (**Fig. 4A-C**). Following the TGD phase, two surviving mice which showed complete tumor regression (CR) from the SB 11285 group, four survivors from the CP group with CRs, and nine survivors from the combination treatment with CRs were subjected to tumor rechallenge studies. All surviving animals in the treated groups completely rejected the tumor growth monitored to 43 days whereas all animals (n = 8) in the control group did not survive by day 37. These observations suggested that the I.T. dosing of SB 11285 alone and in combination with CP, produced durable immune memory in these mice (**Fig. 4D**).

### Dose-response studies of IT., administered SB 11285

In a dose-response study, SB 11285 was tested in the CT-26 colorectal carcinoma syngeneic mouse model at I.T., doses of 2.6 mg/kg (50 μg/mouse), 5.3 mg / kg (100 μg/mouse), and 10.6 mg/kg (200 μg/mouse) dosed on days 0, 1, 3, and 5. There was complete regression (CR) in all 10 mice dosed at the 5.3 and 10.6 mg/kg doses and in 9 out of 10 mice dosed at the 2.6 mg/kg dose of SB 11285 until day 119, the last data point in the study. All the mice tolerated the dosing of SB 11285 well throughout the experiment. (data not shown)

Furthermore, the implant of CT-26 cells into the opposite flank of mice that had no measurable tumor after being previously dosed with SB 11285 resulted in tumor regressions with CRs in 7 of 9 mice previously dosed with 2.6 mg/kg, regressions with CRs in 10 of 10 mice dosed with 5.3 mg/kg, and regressions with CRs in 7 of 10 mice dosed with 10.6 mg/kg. All naïve mice implanted with the same CT-26 cells at the same time developed growing tumors. This observation suggests that the I.T. dosing of SB 11285 produced immune memory in these mice. The immune memory and tumor regression was maximal for mice treated with the 5.3 mg/kg dose.

### The induction of immune memory by SB 11285 in tumor models

At the end of the efficacy study, the complete responders were re-challenged with A20 cells implanted subcutaneously on the opposite flank (**Fig. 4A, B**). Female BALB/c mice were eighteen or twenty-five weeks old (naïve controls and complete regression responder animals from the A20 lymphoma study, respectively) on Day 1 of the study and had a body weight (BW) range of 17.0 to 26.5 g. No treatments were administered in any groups. Group 1 served as the growth control and consisted of eight naïve BALB/c female mice, Group 2 had two survivors from the SB 11285 treatment, Group 3 had four survivors from cyclophosphamide treatment, Group 4 consisted of nine survivors from cyclophosphamide, in combination with SB 11285 treatment. The animals were monitored, and tumor volumes were measured for 38 days post-implantation. All animals in Groups 2, 3 and 4, showed the absence of tumors when monitored up to Day 38, thus showing the induction of immune memory in these animals. On day 38, control animals had MTV of 1667 ± 510 mm^3^.

I.T., dosing of SB 11285 in the B16F10 melanoma model also resulted in no measurable tumor with CRs in 7 of 10 mice dosed at 10.6 mg/kg, in 6 of 10 mice dosed at 5.3 mg/kg, and in 5 of 10 mice dosed at 2.6 mg / kg with durability of response observed until day 113, the last day of the experiment. Furthermore, the implant of B16F10 into the opposite flank of mice, that had no measurable tumor after SB 11285 dosing, resulted in CRs in 2 to 4 mice in each treatment group with no measurable tumor on the opposite flank with durability of response observed till day 113, the last day of the experiment. All naïve mice implanted with the same B16F10 cells at the same time developed growing tumors by day 5. These observations suggest that the I.T., dosing of SB 11285 produced an immune memory in C57BL/6 tumor-bearing mice; however, the extent and persistence of the immune memory was heterogeneous in this model (data not shown).

### Combination Studies of SB 11285 With Checkpoint Inhibitors Demonstrate Profound Anti-Tumor Effects

The efficacy of SB 11285 in combination with, anti-CTLA-4 antibody, a Checkpoint inhibitor, was assessed in the CT26 model. Following the establishment of the subcutaneous tumors with a mean tumor volume (MTV) of 100 mm^3^, I.T., administration of SB 11285 alone, and in combination with the anti-CTLA-4 antibody was initiated at 50 μg/animal of SB 11285 on days 1, 4, 7, 10, and 17, and with anti-CTLA-4 administered I.P., at 5 mg/kg on Day 1 and at 2.5 mg/kg on Days 4 and 7. Highly significant TGD (p<0.001) was observed with 92%, and 177% for SB 11285 and its combination with anti-CTLA-4 antibody respectively with 17% for the anti-CTLA4 alone group, which was not significant compared to the vehicle group. (**Fig. S13)**.

### I.T., SB 11285 Shows Potent Abscopal Effects

We evaluated the abscopal effect of I.T., administered SB 11285 in the CT26 murine colon carcinoma model using female BALB/c mice bearing tumors on both flanks. The I.T., injection of SB 11285 into one flank in BALB/c mice bearing bilateral CT26 tumors demonstrated significant regression of the contra-lateral tumors which is consistent with the induction of adaptive immunity resulting from the administration of SB 11285 to produce antitumor immunity in distal tumors **(Fig. 5)**.

**Fig. 5.**
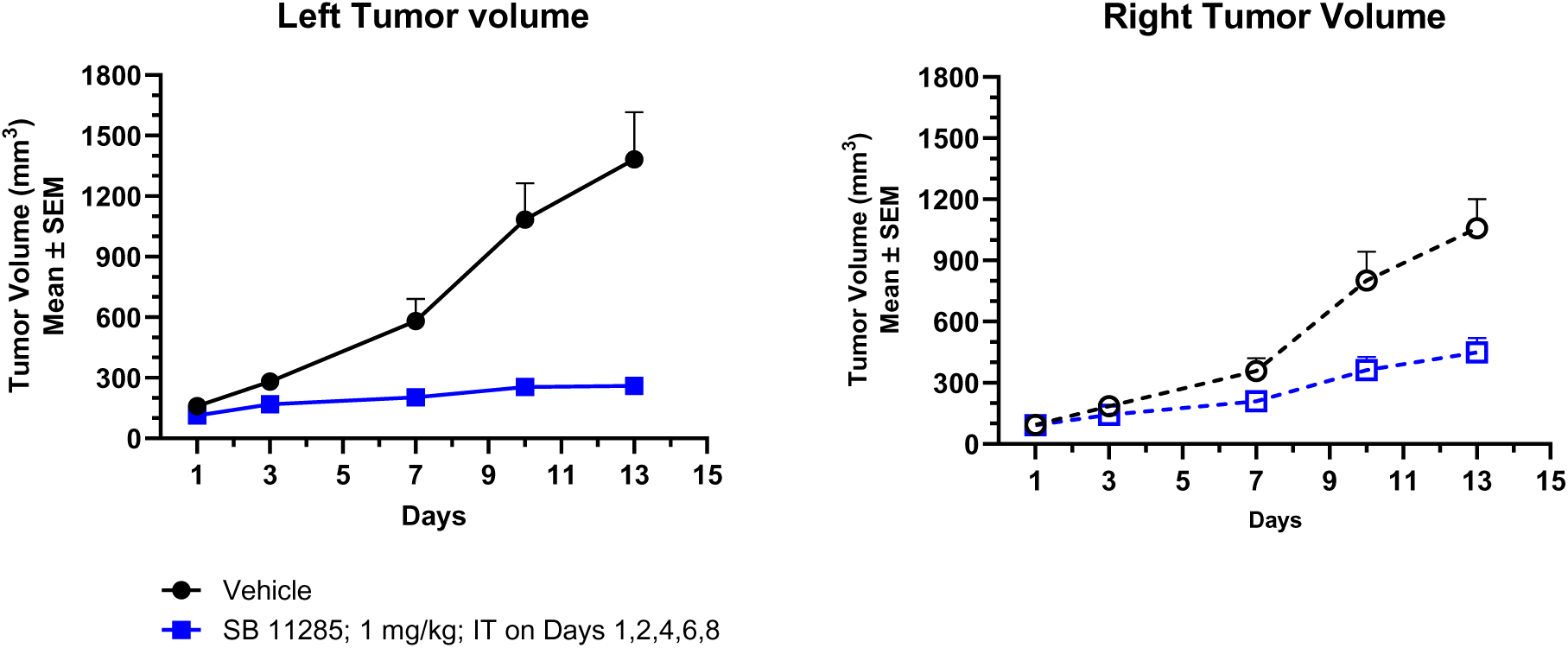
Abscopal antitumoral activity of SB 11285 when administered intratumorally in CT26 colorectal carcinoma model. Groups of female BALB/c mice (n = 10) were implanted with CT26 tumor cells on both right and left flanks. The animals were sorted into two groups (n=10/group) based on tumor volume and administered SB 11285 I.T., at 5 mg/kg (100 μg/mouse) or saline on the left flank on days 1, 2, 4, 6, and 8. Tumors were measured twice weekly for the duration of the study and tumor size was calculated.

### Systemic Administration of SB 11285 Induces Profound Anti-Tumor Efficacy and Immune Memory

A serious limitation of the I.T. treatment approach in cancer is the limited number tumors of cancers that can be treated, with the necessity for specialized expertise and resources for I.T. injections. For STING-based immunotherapy to be more broadly accessible to patients, a systemically bioavailable STING agonist that can be safely administered is highly desirable which could facilitate trafficking of the activated CD8+ T cells from the periphery into the tumor site for effective immunotherapy and could facilitate the conversion of “cold” tumors into “hot tumors” (56). We therefore considered the evaluation of systemically administered SB 11285 analogs.

Indeed, in the CT26 model, intraperitoneal (I.P.) administration of SB 11285 at 1, 3, and 9 mg/kg (four injections each dose) resulted in dose-dependent anti-tumor activity with tumor growth inhibition compared to vehicle control. The I.P., administered SB 11285 also potentiated the antitumor activity of anti-CTLA antibody r. Interestingly, the intravenous (I.V.) administration of SB 11285 at 9 mg/kg alone showed more potent antitumor activity with a TGI of 95% compared to I.P., dosing at 9 mg/kg, with TGD being 81% vs. 49% for I.V., vs I.P. (**Fig. 6**).

**Fig. 6.**
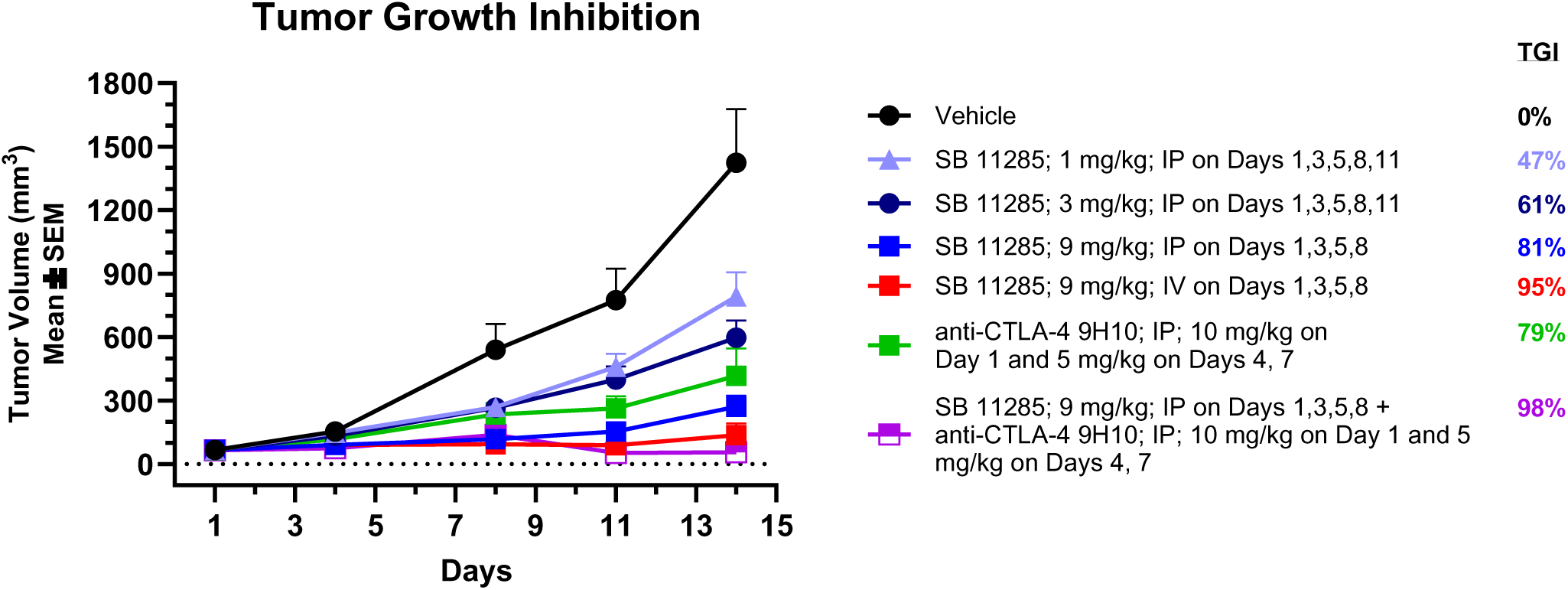
Anti-tumor efficacy of I.V., and I.P., administered SB 11285 alone and in combination with anti-CTLA Ab in CT26 mouse colorectal carcinoma model. Groups of female BALB/c mice (n = 10) were implanted with CT26 tumor cells on the right flank. When the tumors grew to an average MTV of 100 mm^3^, two of the treatment groups were administered I.P., SB 11285 at 1, 3 or 9 mg/kg or I.V., SB 11285 at 9 mg/kg on days 1, 3, 5, and 8. Additional treatment groups received I. P., anti-CTLA antibody at 5 mg/kg on days 1, 4, and 7 either alone or in combination with I.P. SB 11285 at 9 mg/kg treatment group. Tumor volumes and TGIs for each of the treatment groups were assessed on days as indicated.

In a dose-ranging study, the I.V., administration of SB 11285 in the CT-26 model resulted in no measurable tumors with CRs in 1 out of 10 mice dosed at the 3.2 mg/kg dose and CRs in 5 of 10 mice dosed at the 5.3 mg/kg doses (four injections each dose). The CRs observed in these mice were durable and the tumors did not grow back till the last day of the experiment on day 119. Additionally, the implant of CT-26 into the opposite flank of mice with CRs after SB 11285 dosing, resulted in all mice in each treatment group (5 of 5 from 5.3 mg/kg and 1 from 3.2 mg /kg) displaying CRs on the opposite flank. All naïve mice implanted with the same cells at the same time developed growing tumors. These observations suggested that the intravenous dosing of SB 11285 produced an immune memory in these mice (data not shown).

### I.P., and I.V., SB 11285 Induces Profound Anti-tumor Efficacy in Orthotopic Tumor Models

Given the potent antitumor activity of systemically administered SB 11285 in subcutaneous tumor models, we next evaluated the efficacy of SB 11285 in syngeneic orthotopic tumor models. Tumors of orthotopic models grow in the organ of origin. In these models, there is considerable interaction with stromal cells, as well as tissue-specific immune cells that allow tumor growth mimicking the original organ tissue (62). The interaction of tumor cells with tissue-specific stroma cells not only affects tumor growth and differentiation, but also efficacy of a test agent. Moreover, orthotopic tumors can metastasize with specificities comparable to human cancers. Furthermore, the use of fluorescently labeled tumor cells for implantation, facilitates imaging of the tumors therefore providing knowledge about the growth characteristics of the tumors, as well as tumor metastasis (62). In this regard, the 4T1 mammary carcinoma is a transplantable murine tumor cell line that is highly tumorigenic, invasive, and spontaneously metastasizes from the primary tumor in the mammary gland to multiple distant sites. The metastatic spread of 4T1 tumors to the draining lymph nodes and other organs is very similar to that of human mammary cancer and therefore the 4T1 model is considered to have high clinical translation (62).

We evaluated SB 11285 by I.P., administration in the 4T1 syngeneic mouse tumor model following implantation with 4T1-luc2 cells that enabled imaging and visualization of tumor metastasis. Administration of SB 11285 at 10 mg/kg on days 5, 7, 9, 11, 13, 15, 17, and 19 resulted in a significant decrease in the MTV of the primary tumor **(Fig. 7)**. The compound was well tolerated, and significant inhibition of tumor metastasis was also noted on Day 19 by the whole-body imaging of mice treated with SB 11285 compared to untreated mice (**Fig. 7**) with several treated mice showing no metastases.

**Fig. 7.**
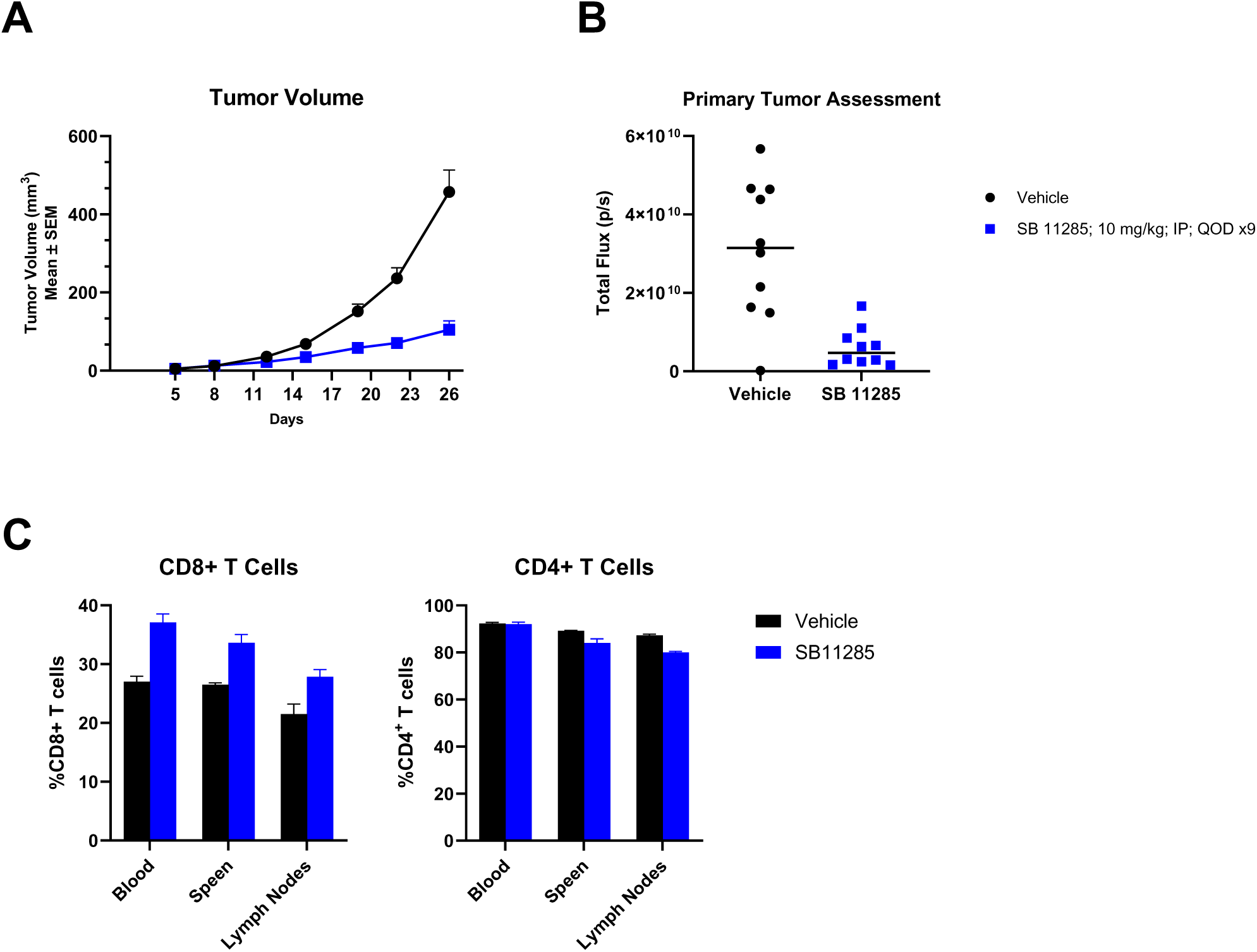
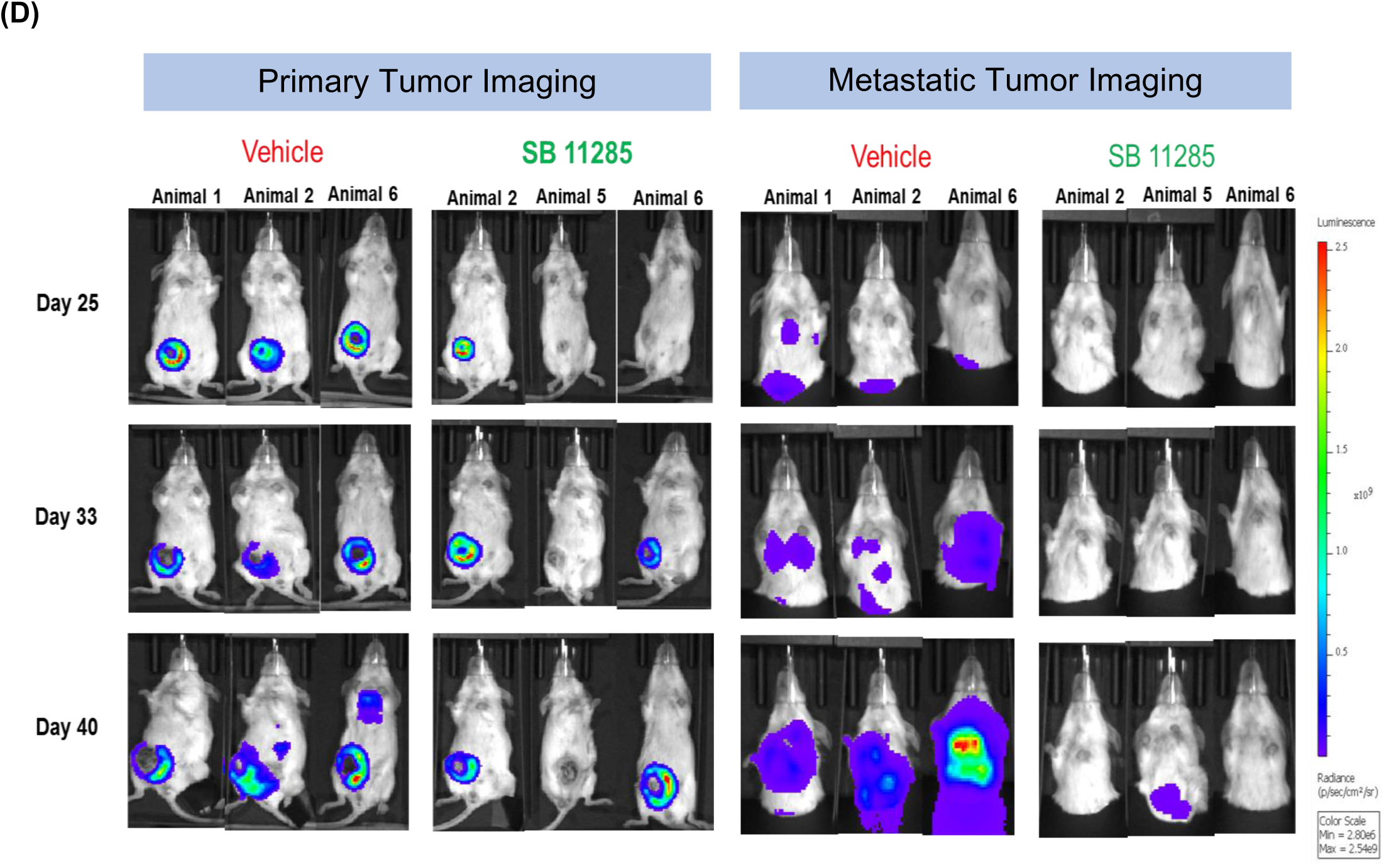
Anti-tumor activity of I.P., SB 11285 in the 4T1 breast cancer model. Groups of nine-week-old female BALB/c mice were implanted with 1 x 10^5^ 4T1-luc2 cells in PBS (0.1 mL) in the mammary fat pad and the animals were sorted into four groups each consisting of either five mice (for sampling) or ten mice (for efficacy). SB 11285 was administered I.P., at 10 mg/kg on Days 5, 7, 9, 11, 13, 15, 17, and 19 in sterile saline. **(A)** Primary tumor measurements were performed on indicated days and MTV was plotted until day 26. (**B)**. Whole body image total flux was plotted as indicated. (**C). Flow cytometric analysis**. Blood, spleen, and lymph nodes were collected on day 19 to analyse T cell populations by flow cytometry. Spleen and lymph node samples were homogenized using RPMI-1640 media and the red blood cells lysed with ACK buffer prior to analysis. Isotype control antibodies were used as negative staining controls. For staining of internal markers, cells were permeabilized with cytofix/perm buffer. Internal marker staining was carried out using 0.1 μg of each antibody diluted in 100 μL of PB. For surface staining of cells, single cell suspensions from tumor, spleen and lymph nodes were seeded on to 96 well U bottom plates. 50 μL of different antibodies were added to the respective wells for viability staining. Plates were placed in BD FACSVerse flow cytometer with appropriate compensation controls and data analysis was carried out using FlowJo software. Data was plotted as the % of CD8+ T cells and the % of CD4+ T cells in spleen, lymph nodes and blood. **D**. **Inhibition of tumor metastasis.** Whole body imaging was carried out to assess Luciferase activity in live animals using IVIS® SpectrumCT equipped with a CCD camera, mounted on a light-tight specimen chamber. On the day of imaging, animals were injected with 0.22 μm-filter sterilized VivoGlo™ D-Luciferin substrate (150 mg/kg, I.P.) and placed in anesthesia induction chamber. Upon sedation, animals were placed in a ventral position in the imaging chamber, equipped with stage-heated physiological temperature, for image acquisition ten minutes following luciferin substrate injection. Animals were imaged with the primary tumor unshielded followed by shielded imaging, and the images were used for determination of the primary tumor and metastasis flux values were quantified and reported as 10^6^ photons per second (p/s). The light emitted from the bioluminescent cells was detected, digitalized, and electronically displayed as a pseudo-color overlay onto a gray-scale photographic image acquired immediately prior to bioluminescent imaging, to allow for the anatomical localization of the signal. Data were analyzed and exported using Living Image software. Representative images of animals are shown.

The percentage of CD8+ T cells, CD4+ T cells, and MDSCs in spleen, lymph nodes and blood were also measured by flow cytometry on day 19, after treatment initiation. The induction of CD8+ T cells were noted in blood, lymph nodes and spleen, whereas there was a decrease CD4+ T cells and gMDSCs in lymph nodes and spleen. It was interesting to note that there was an increase in the percentage of mMDSCs in the lymph node with no corresponding changes in blood and spleen (**Fig. 7**).

Next, we evaluated the anti-tumor activity of SB 11285 in an orthotopic rat bladder cancer model. Bladder tumors were developed by implanting Wistar rat bladder tumor (NBT-II) cells into the bladders of syngeneic immunocompetent rats. The anti-tumor activity of SB 11285 was then evaluated at I.V., doses of 0.5, 1.5 and 3 mg/kg, and Docetaxel 7.5 mg/kg I.V., as a measure of decrease in bladder weight in treated vs. control animals following necropsy on day 31 after initiation of treatment. The mean bladder weights in all the treatment groups were significantly lower than the vehicle control group which had tumor burden (**Fig. 8**). A gross pathological evaluation showed 3 to 5 focal tumor nodules around the bladder in vehicle control group animals but no visible macroscopic nodules in any of the treatment groups (**Fig. 8**).

**Fig. 8.**
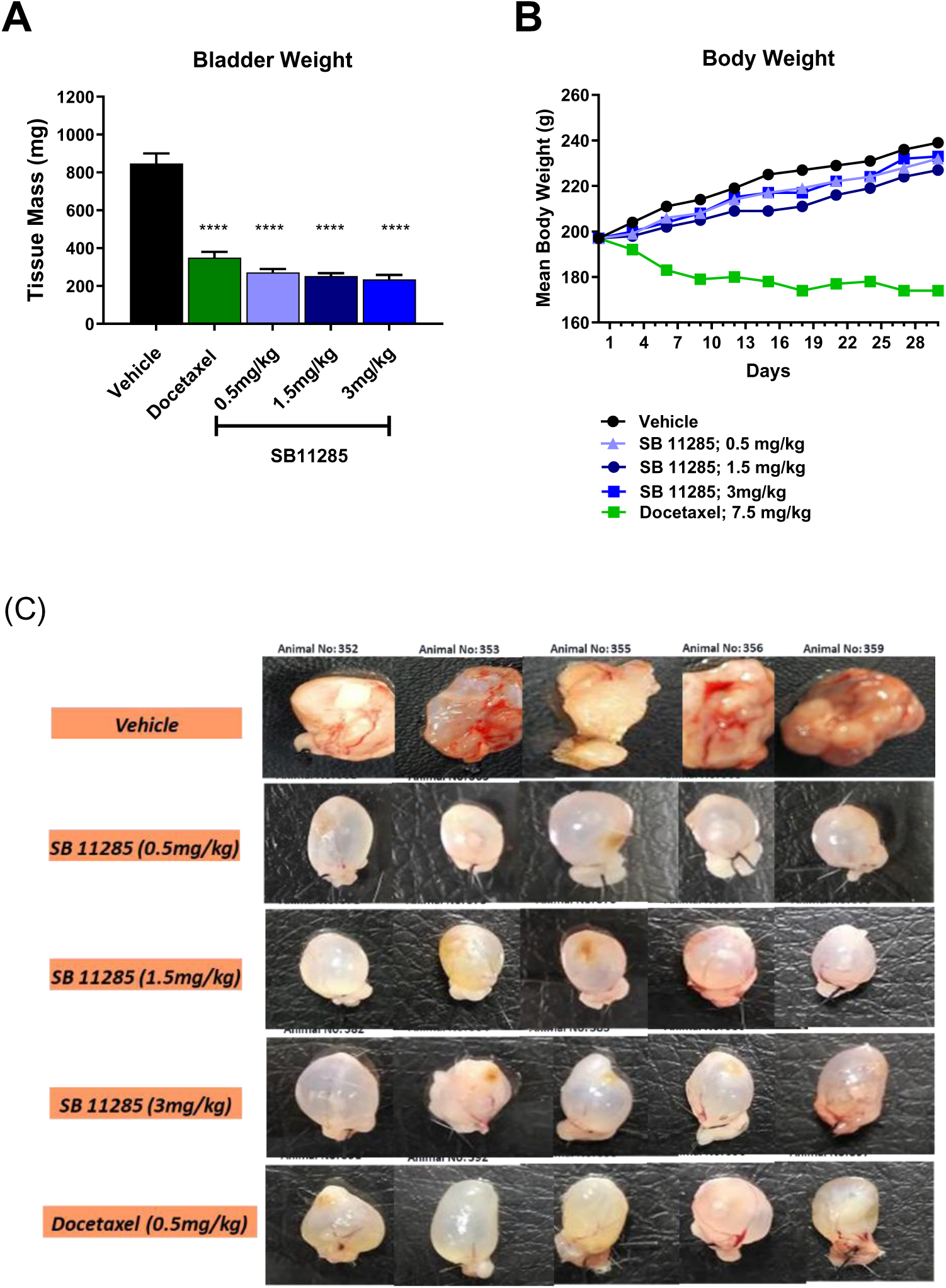
Evaluation of I.V. SB 11285 in the NBT II rat bladder cancer model. Syngeneic Rat NBT II Bladder Cancer Model. Rats (n = 10) were anesthetized, and a small incision was made on the lower abdomen region and NBT-II cells (2 x 10^6^ in 50 µL of PBS) were injected slowly in the bladder wall. The bladder dome was placed back into the abdomen and the site of incision was sutured closed. After 6 days, animals were randomized based on their body weights, and SB 11285 or docetaxel were administered indicated. **(A).** Mean body weights were recorded throughout the study period and plotted as shown. **(B).** At the end of 31 days, animals were euthanized, bladder removed, trimmed of fascia, and the bladder weights were plotted as shown. **(C).** Bladders were photographed and gross pathology evaluation was conducted to reveal tumor nodules; representative images are shown. The statistical significance of tumor inhibition was performed by One-way ANOVA followed by Dunnett’s t-test using Graph Pad Prism *p* values <0.05 was considered statistically significant between the groups with “****”) at *P* ≤ 0.0001 considered extremely significant.

The absence of focal tumor nodules coupled with significantly lower bladder weights in the SB 11285-, and Docetaxel-treated groups compared to vehicle control group demonstrated the potent anti-tumor activity of SB 11285, and Docetaxel. However, treatment with Docetaxel showed moderate to severe body weight loss (mean weight loss of 12 %) during the study period (**Fig. 8)**, whereas SB 11285-treated animals showed an increase in body weights. Furthermore, the I.V., administration of SB 11285 was well tolerated with no mortality or body weight loss and with no visible clinical signs or abnormal behaviour observed in any of the treatment groups. Also, there were no distant metastases observed in any of the treatment groups.

### I.V., SB 11285 Enhances the Anti-tumor Activity of Anti-PD1 Antibody

The I.V. administered SB 11285 at 3 mg/kg was also evaluated in the MC-38 colon carcinoma mouse model in combination with anti-PD1 antibody. While SB 11285 alone showed 4 CRs out of the 8 treated animals, there were no CRs in any of the PD-1 treated mice. A combination of SB 11285 with anti-PD-1 antibody gave 7 CRs out of 8 treated animals thereby potently enhancing the weak antitumor activity of anti-PD1 antibody **(Fig. 9)**.

**Fig. 9.**
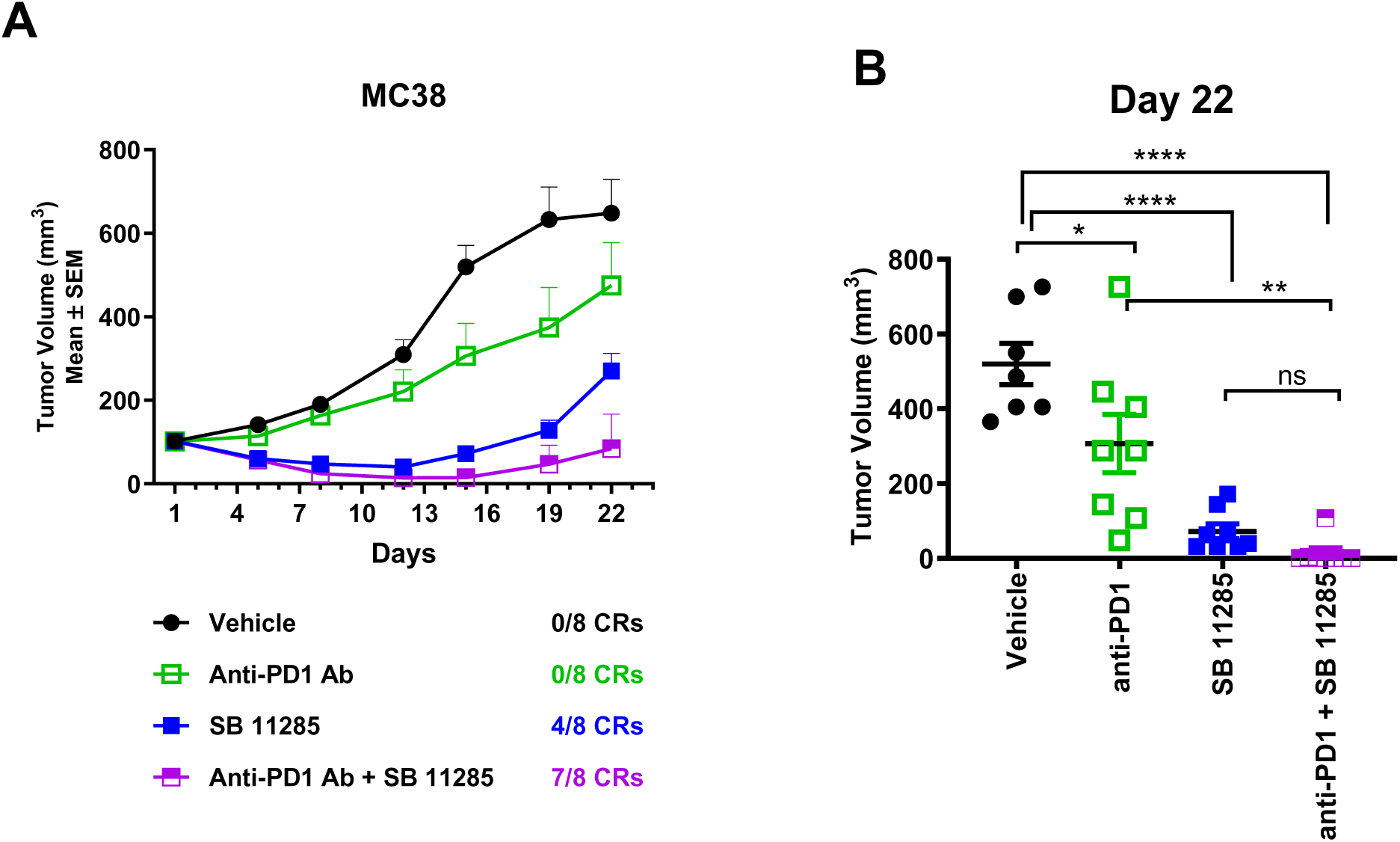
Combination study of I.V., SB 11285 with anti-PD1 antibody in the MC-38 colon carcinoma model. **(A).** Groups of female C57BL/6 mice (n = 8) were injected with 5×10^5^ MC38 tumor cells in 0% Matrigel S.C., in the right flank at a cell Injection volume of 0.1 mL/mouse. Once the tumor volumes reached around 80 to 120 mm^3^, the animals were randomized into groups either to receive treatment with the vehicle or the test compounds. The anti-PD-1 RMP1-14 antibody was administered I.P., biweekly at 5 mg/kg. SB 11285 was administered I.V., at 3 mg/kg on days 1, 5, 9, and 14 alone or in combination with the anti-PD1 antibody. Animals were monitored as a group until day 24. MTVs were measured on indicated days and plotted as shown. **(B).** Mouse tumor volumes plotted as mean ± SEM, on day 22 following the treatment with vehicle, anti-PD1 antibody, SB 11285, or SB 11285 + anti-PD1 antibody. Statistical significance was assessed by unpaired t-test, two tailed.

Thus, our studies indicate that SB 11285 can augment the efficacy of both anti-PD1 and anti-CTLA antibodies.

## Discussion

In this study, we investigated a new class of purine-pyrimidine cyclic dinucleotides with 3’,5’, 2’,3’ and 3’,2’ internucleotidic connectivities as STING agonists. We carried out molecular modeling of 3’,5’, Pu-Py CDNs based on the crystal structure of CDNs with hSTING protein and designed a focused library of compounds to identify leads using cell-based reporter assays. Following extensive structure-activity relationship studies for optimizing hSTING agonism, we found that the chemo-selective attachment of an ar-alkyl chain of different lengths to the single phosphorothioate backbone of a PuPy-CDN influenced the physicochemical attributes of the CDN resulting in significant enhancement of its lipophilicity.

Furthermore, the lead compound SB 11285, was found to remarkably self-assemble into nano structures in aqueous media as assessed by Scanning Electron Microscopy (data not shown). SB 11285 has been established as a highly specific STING agonist. Unlike the natural STING agonist cGAMP, and the synthetic STING agonist ADU-S100, SB 11285 was taken up by multiple cell types without the aid of transfection reagents and produced robust hSTING-dependent or mSTING-dependent induction of IRF and NF-κB signaling pathways. SB 11285 was found to be significantly more potent than both cGAMP, and the synthetic STING agonist ADU-S100 which are both 2’,3’-purine-purine-purine-based CDNs.

In this study, the binding modes of the 3’,5’-PuPy-CDNs with hSTING, have been computationally modeled. Thus, while interactions with key residues Tyr167 and Arg238 within the CDN pocket facilitated the binding of SB 11285 to hSTING, additional interactions between the aryl-alkyl chain of SB 11285 with the amino acid residues from the gatekeeper loop of B-chain dimer enhanced the stabilization of STING dimer interface and tighter binding of SB 11285. Based on our MD simulations of STING CTT structure, it is expected that the ar-alkyl chain of SB 11285 may also potentially interact with the C-terminal tail (CTT) of STING. These additional interactions between SB 11285 functionalities and amino acid residues in STING may help to potently activate STING as supported by various studies reported here. SB 11285 was also found to potently activate multiple hSTING polymorphic variants.

In in vitro studies, SB 11285 was found to dose-dependently induce type I, type II IFNs and type III, and other immune mediators (IL-12p70, IL-18, GM-CSF etc.), pro-and anti-inflammatory cytokines, and various chemokines in PBMCs from all donors, in a profile similar to that produced by cGAMP and ADU-S100 both of which were however less potent compared to SB 11285. A preponderance of the in vitro data shows that SB 11285 induces IFN-β production in APCs via the classical STING-TBK1-IRF3 signaling pathway.

In vivo, in syngeneic mouse tumor models, SB 11285 induced Type I and Type II IFNs, and other cytokines and chemokines. Additionally, the pharmacodynamic studies of SB 11285 in syngeneic mouse tumor models showed the induction of CD8 +T cells and reduction of Tregs, besides the induction of IFNs and important antitumor cytokines, consistent with its antitumor effects. Importantly, the administration of SB 11285 by I.T., I.V., or I.P., routes showed potent anti-tumor activity in multiple subcutaneous and orthotopic syngeneic mouse and rat tumor models, that was also associated with complete tumor regression, the induction of immune memory and abscopal effects. The mechanism of the anti-tumor effect observed with the compounds tested was consistent with STING activation, with the anti-tumor effect being dependent upon T cells, specifically CD8+ T cells (58, 59). Furthermore, immunohistochemistry of cells from the TME, from tumor-bearing mice treated with SB 11285, also indicated the induction of activated macrophages and NK cells, in addition to CD8+ T cells, that could collectively amplify and promote the observed potent anti-tumor activity.

Overall, we hypothesize that the activation of the STING pathway by SB 11285 in the circulating immune cells or tumor-resident immune cells that carry tumor antigens will enable the efficient processing of the antigens for presentation to naïve T cells and subsequent activation and clonal expansion of tumor-specific CD8+ T cells. It is to be noted that in addition to the T-cell-dependent component of the therapeutic effect of STING agonists in vivo, additional cytokines and chemokines induced by SB 11285 may also have beneficial antitumor effects. Thus, IFN expression can promote apoptosis of tumor cells, and the induction of TNF-alpha may cause tumor-associated vascular destruction and tumor killing. Indeed, in vitro, the treatment of tumor cells with SB 11285 caused the induction of IFN-mediated apoptosis of tumor cells. Furthermore, the induction of chemokines by SB 11285 may contribute to the effective migration of activated T cells into the TME at the tumor site. It appears likely that the induction of cytokines and chemokines by SB 11285 may also contribute to effective interplay between innate and adaptive immunity. Overall, this multi-faceted mechanism of action involving apoptosis, and induction of innate and adaptive immune response may explain the potent antitumor effects of systemically and intratumorally administered SB 11285 against a range of cancers in subcutaneous and orthotopic murine tumor models. However, a detailed analysis of the subsets of immune cells in the TME and the evaluation of whether a reprogramming of the TME occurs that promotes efficient tumor killing, is needed to fully understand the anti-tumor activity profile of SB 11285.

It is pertinent to mention the combination of SB 11285 with the checkpoint inhibitors, anti-CTLA-4 and anti-PD1 antibodies, or cytotoxic agents such as cyclophosphamide showed enhanced antitumor efficacy in different tumor models. Recently, it was also reported that SB 11285 in combination with low dose radiation therapy showed potent additive anti-tumor effects in the head and neck cancer model of mice bearing FaDu or Detroit562 tumors (65). This study revealed that STING expression in the tumor is required for maximal therapeutic effects induced by radiation. Thus, SB 11285 can be productively combined with different classes of antitumor agents and treatment strategies for enhanced antitumor activity against different cancers.

In conclusion, our preclinical studies demonstrate that the clinical candidate SB 11285 has high translational potential as an immunotherapeutic agent for durable anti-tumor efficacy in multiple tumor types. Our studies also demonstrate that an expanded repertoire of cell-permeable 3’,5’, 2’,3’, and 3’,2’-PuPy-CDN compounds can be developed as STING agonists (**Fig. 1**) that can be administered by multiple routes for therapeutic application as antivirals, and antitumor agents, as well as, vaccine adjuvants. Based on our results, it may be possible to activate the STING pathway potently and safely for anti-tumor effects with appropriately designed canonical Pu-Py-3,5’-CDNs as exemplified by SB 11285, a potent STING agonist for administration by multiple routes.

The I.V. administered SB 11285 is currently under evaluation in human clinical trials in multiple tumor types as the first systemically administered 3’,5’-CDN, alone and in combination with checkpoint inhibitors (66, 67).

## Acknowledgements

We thank Leena Suppiah, Dr. Diane Schmidt, Dr. Lakshmi Bhagat, and other members of the Spring Bank Discovery team for helpful discussions and excellent technical assistance. We also thank Dr. Sarah Batey and Dr. Kin-Mei Leung of InvoX Pharma Ltd., for their insightful review of the manuscript. We thank Charles River laboratories and Syngene International and other CROs for their excellent contributions in the evaluation of our compounds in different tumor models. We also acknowledge the assistance of the University of Massachusetts Medical Center, MA for providing access to the Core facilities in obtaining SEM images, Microscopy Imaging studies, and Immunohistochemistry of tissue samples.

