## Supplemental tables and Figures for "SB 11285, a Novel Canonical Purine-Pyrimidine Cyclic Dinucleotide, is a Systemically Bioavailable STING Agonist for Cancer Immunotherapy"

### Summary

The stimulator of interferon genes (STING) pathway of cytosolic DNA sensing is recognized as an important innate immune sensing mechanism and the discovery of compounds that activate STING has emerged as an exciting area for the advancement of novel immunotherapeutics against cancer. We describe here the discovery and development of a new class of a canonical 3',5' purine-pyrimidine cyclic dinucleotides (PuPy-CDNs) that potently activate the STING pathway for antitumor activity. The lead candidate SB 11285 which is readily cell permeable and is substantially more potent than the known non-canonical 2,3'-CDNs. Furthermore, SB 11285 induced profound antitumor activity when administered by intratumoral, intravenous and intraperitoneal routes in subcutaneous and orthotopic models of syngeneic tumor models, inducing complete tumor regression of established tumors, generating substantial adaptive immune responses, and providing abscopal antitumor effects, as well as, long-lived immunologic memory. When used in combination with checkpoint inhibitors, or cytotoxic agents, SB 11285 showed enhanced antitumor efficacy. The anti-tumor effects were dependent upon the induction of CD8+ T cells, as well as activated macrophages and NK cells in the TME, that could collectively amplify and promote profound anti-tumor response. Our studies demonstrate that the STING pathway can be potently and safely activated for anti-tumor activity using functionalized canonical 3',5'-purine-pyrimidine class of CDNs as STING agonists exemplified by SB 11285 for administration by multiple routes.

The I.V. administered SB 11285 is currently under evaluation as the first systemically administered CDN in human clinical trials in multiple tumor types.

**Note**: In all studies reported in Tables, the mean values ± SD are from three independent experiments.

Fold-change is expressed relative to the no compound control.

**Table S1. Activity of SB 11285 in human THP-1 cells does not require cGAS or IFI16.**

| **Compound** | **Human THP-1 cell line – IRF-3 reporter activation** | | | | | |
|---|---|---|---|---|---|---|
| | **Endogenous STING-HAQ** | | **KO-cGAS** | | **KO-IFI16** | |
| | **$EC_{50}$ (μM)** | **Maximum Fold-change** | **$EC_{50}$ (μM)** | **Maximum Fold change** | **$EC_{50}$ (μM)** | **Maximum Fold change** |
| **SB 11285** | 0.14 ± 0.01 | 89 ± 27 | 0.15 ± 0.01 | 242 ± 117 | 0.15 ± 0.01 | 84 ± 31 |

**Table S2. Activity of 10 μM SB 11285 in HEK293 cells stably expressing individual human or mouse innate immune receptors.**

| HEK293 cell line | | Fold-change | |
|---|---|---|---|
| | | SB 11285 | Positive control |
| Toll-like receptors(TLRs) | Human TLR2 | 1 ± 0 | 36 ± 2 |
| | Mouse TLR2 | 1 ± 0 | 25 ± 1 |
| | Human TLR3 | 1 ± 0 | 14 ± 0 |
| | Mouse TLR3 | 2 ± 0 | 25 ± 1 |
| | Human TLR4 | 1 ± 0 | 15 ± 1 |
| | Mouse TLR4 | 1 ± 0 | 14 ± 1 |
| | Human TLR5 | 1 ± 0 | 27 ± 0 |
| | Mouse TLR5 | 1 ± 0 | 35 ± 1 |
| | Human TLR7 | 3 ± 0 | 34 ± 1 |
| | Mouse TLR7 | 1 ± 0 | 18 ± 0 |
| | Human TLR8 | 1 ± 0 | 15 ± 1 |
| | Mouse TLR8 | 1 ± 0 | 15 ± 1 |
| | Human TLR9 | 1 ± 0 | 10 ± 0 |
| | Mouse TLR9 | 1 ± 0 | 18 ± 1 |
| | Mouse TLR13 | 1 ± 0 | 41 ± 0 |
| Nod-like receptors(NLRs) | Human NOD1 | 2 ± 0 | 14 ± 1 |
| | Mouse NOD1 | 2 ± 0 | 13 ± 0 |
| | Human NOD2 | 1 ± 0 | 15 ± 1 |
| | Mouse NOD2 | 1 ± 0 | 6 ± 1 |
| C-type lectin receptors (CLRs) | Human Dectin1a | 1 ± 0 | 13 ± 0 |
| | Mouse Dectin1a | 1 ± 0 | 12 ± 1 |
| | Human Dectin1b | 2 ± 0 | 17 ± 1 |
| | Human Mincle | 1 ± 0 | 16 ± 0 |
| Control TLR$^-$/NLR$^-$/CLR$^-$ | Control cell line 1 | 2 ± 0 | 17 ± 1 |
| | Control cell line 2 | 2 ± 0 | 35 ± 1 |
| | Control cell line 3 | 1 ± 0 | 29 ± 2 |
| | Control cell line 4 | 1 ± 0 | 11 ± 0 |
| | Control cell line 5 | 1 ± 0 | 19 ± 1 |
| RIG-I-like receptors(RLRs) | Human RIG-I | 8 ± 1 | 30 ± 2 |
| | Human MDA5 | 12 ± 0 | 8 ± 0 |
| | Control RIG-I$^-$/MDA5$^-$ | 10 ± 0 | 84 ± 14 |

**Table S3. The Activation of Mouse STING by SB 11285**

| Compound | Mouse B16 cell line – IRF3 reporter activation | | | |
|---|---|---|---|---|
| | Endogenous STING | | KO-STING | |
| | $EC_{50}$ (μM) | Maximum Fold-change | $EC_{50}$ (μM) | Maximum Fold change |
| **SB 11285** | 0.62 ± 0.04 | 2.7 ± 0.3 | >10 | 1.1 ± 0.3 |
| **cGAMP** | >10 | 1.9 ± 0.5 | >10 | 1.2 ± 0.4 |
| **ADU-S100** | 2.1 ± 0.2 | 2.6 ± 0.3 | >10 | 1.2 ± 0.2 |

**Table S4. The induction of cytokines by SB 11285 in murine B16 splenocytes.**

Murine splenocytes from healthy CD-1 mice (n = 4 animals) were stimulated with SB 11285, cGAMP and ADU-S100. Cytokine responses were measured 24 hours after treatment by multiplex analysis using a 36-plex Luminex assay and ELISA (IFN-β).

| Cytokine | | SB 11285 | | cGAMP | | ADU-S100 | |
|---|---|---|---|---|---|---|---|
| | | $EC_{50}$ (μM) | Max fold change | $EC_{50}$(μM) | Max fold change | $EC_{50}$ (μM) | Max fold change |
| **Type I IFN** | **IFN-α** | 0.2 ± 0.1 | 35 ± 21 | >10 | 4 ± 3 | >10 | 13 ± 5 |
| | **IFN-β** | 0.2 ± 0.1 | 54 ± 26 | >10 | 1 ± 0 | >10 | 27 ± 18 |
| **Immuno-modulatory** | **IL-18** | 0.02 ± 0.02 | 11 ± 12 | >10 | 8 ± 9 | 0.6 ± 0.4 | 13 ± 5 |
| **Pro/Anti inflammatory** | **TNF-α** | 0.04 ± 0.06 | 26 ± 14 | >10 | 9 ± 2 | 1.4 ± 0.5 | 25 ± 11 |
| | **IL-6** | 0.02 ± 0.01 | 94 ± 51 | >10 | 42 ± 19 | 1.7 ± 1.7 | 49 ± 19 |
| | **IL-17A** | 0.003 ± 0.002 | 11 ± 6 | 1.5 ± 0.8 | 9 ± 4 | 0.3 ± 0.2 | 6 ± 2 |
| | **IL-10** | 0.02 ± 0.02 | 9 ± 9 | >10 | 4± 1 | >10 | 6 ± 2 |
| | **IL-31** | 0.006 ± 0.003 | 8 ± 3 | 2 ± 1 | 5 ± 1 | 0.3 ± 0.2 | 5 ± 1 |
| **Chemokines** | **CXCL1** | 0.02 ± 0.01 | 16 ± 5 | >10 | 8 ± 3 | 1.6 ± 0.9 | 13 ± 3 |
| | **IP-10** | 0.03 ± 0.02 | 407 ± 265 | 5.2 ± 3.1 | 200 ± 181 | 2.1 ± 2.9 | 236 ± 186 |
| | **MIP-1α** | 0.006 ± 0.005 | 24 ± 15 | 2.3 ± 0.7 | 19 ± 10 | 0.5 ± 0.4 | 17 ± 9 |
| | **MIP-1β** | 0.005 ± 0.003 | 25 ± 12 | 2.6 ± 1.1 | 25 ± 16 | 0.7 ± 0.3 | 28 ± 13 |
| | **RANTES** | 0.001 ± 0.002 | 18 ± 4 | 0.9 ± 0.3 | 25 ± 12 | 0.1 ± 0.1 | 26 ± 9 |

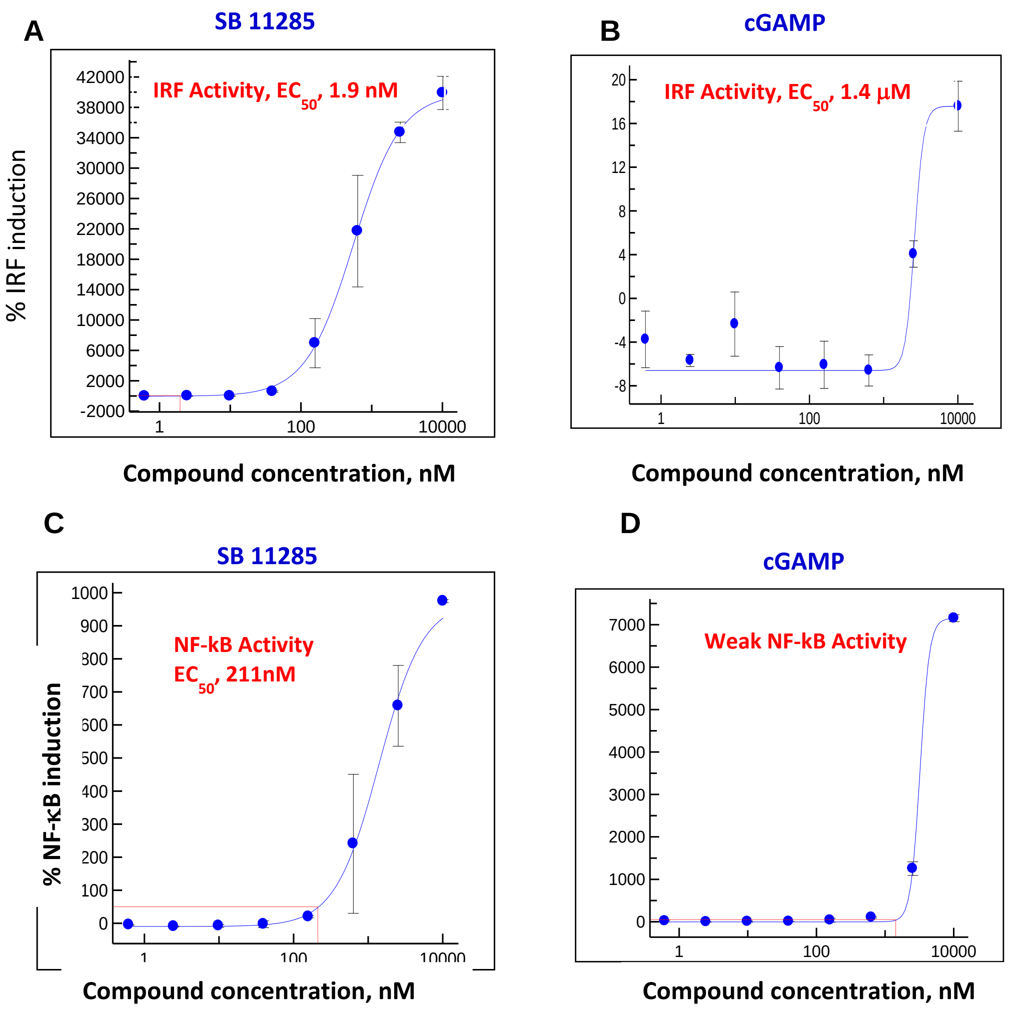

**Fig. S1**. **$EC_{50}$ assessment of SB 11285 and cGAMP using THP-1 cells carrying reporter genes**. THP1 dual cells were treated with various concentrations of SB 11285 or cGAMP or DMSO control with Lipofectamine LTX. Dual cells carry both secreted embryonic alkaline phosphatase (SEAP) reporter gene under the control of an IFN-β minimal promoter fused to five copies of the NF-κB consensus transcriptional response element to measure NF-κB activity and Lucia reporter gene under the control of an ISG54 minimal promoter to measure IRF activity. After 20 hrs., incubation, IRF activity (**A)** for SB 11285 and **(B)** for cGAMP and (**C)** NF–κB activity for SB 11285 and **(D)** for cGAMP was determined by measuring SEAP levels at 620-655 nm. The % Induction was calculated as 5X fold-change in luminescence/absorbance compared to DMSO-treated sample. $EC_{50}$ values were generated by curve fit in Xlfit. IRF activity of SB 11285 was 1.9 nM, while that of cGAMP was 1.4 μM; NF-κB activity of SB 11285 was 211 nM, while that of cGAMP was weak and not calculated.

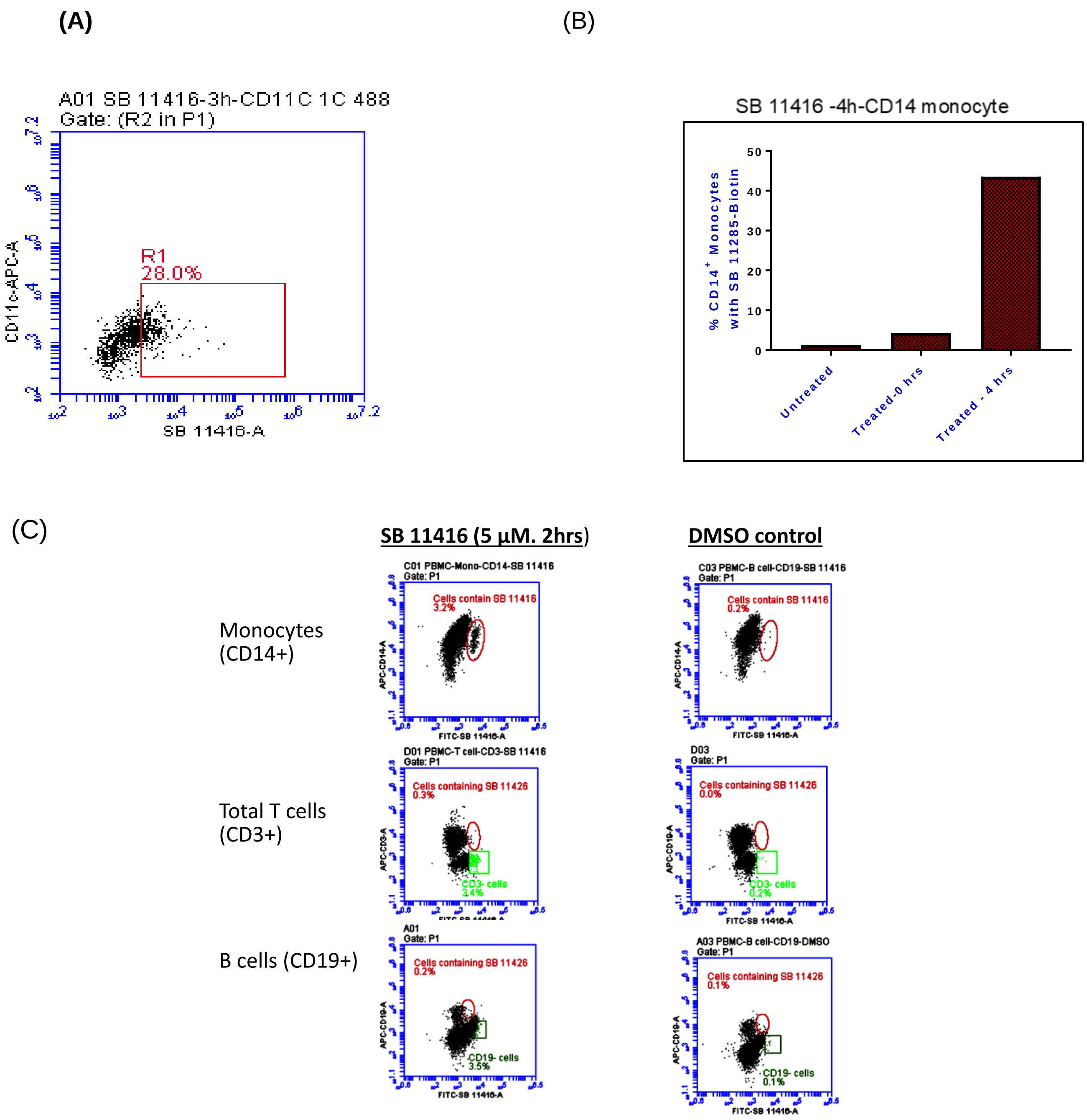

**Fig. S2. Flow cytometric analysis showing significant uptake of SB 11285 with monocytes and dendritic cells. (A).** Human PBMCs were incubated with the biotin-tagged SB 11285 (called SB 11416), alone (5 µM) **(**or DMSO control for 2 hrs. Cells were harvested, fixed and stained with various surface markers such as anti-CD11c+ antibody (dendritic cells) (**B**), anti-CD14 antibody (monocytes) or (**C**) anti-CD3 antibody (total T cells), or anti-CD19 antibody (B cells) followed by staining with Streptavidin-488 for intracellular staining to detect cells with the uptake of SB 11416 and flow cytometric analysis.

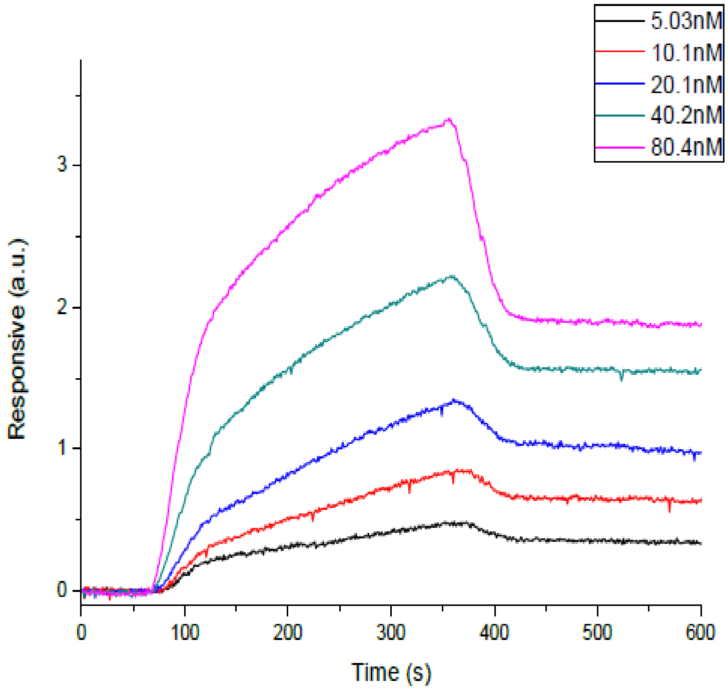

**Fig. S3. Interaction of SB 11285 with STING-CTT by Surface Plasmon Resonance**. Aqueous solutions (0.2 μL) of various concentrations of biotinylated SB 11285 (1mg/mL stock solution) were manually printed onto gold-coated (thickness 47 nm) PlexArray Nanocapture Sensor Chip (Plexera Bioscience, Seattle, WA, US) in replicate at 40% humidity. Each concentration was printed in replicate, and each spot contained 0.2 μL of ligand solution. The chip was incubated in 80% humidity at 4°C for overnight, and rinsed with 10× PBST for 10 min, 1× PBST for 10 min, and deionized water twice for 10 min. The chip was then blocked with 5% (w/v) non-fat milk in water overnight, and washed with 10× PBST for 10 min, 1× PBST for 10 min, and deionized water twice for 10 min before being dried under a stream of nitrogen prior to use. WT-hSTING-CTD (aa138-379) (WT-STING-CTT) was injected into the flow cell. The chip was incubated in 80% humidity at 4°C for overnight, and rinsed with PBST, followed by deionized water. The chip was then blocked with 5% (w/v) non-fat milk in water overnight and washed with PBST followed by deionized water and dried under $N_2$. SPRi measurements were performed with PlexAray HT (Plexera Bioscience, Seattle, WA, US). The signal changes after binding and washing (in AU) were recorded were recorded and analyzed by Data Analysis Module (DAM, Plexera Bioscience, Seattle, WA, US). Kinetic analysis was performed using BIA evaluation 4.1 software (Biacore, Inc.). KD was estimated to be 122 nM.

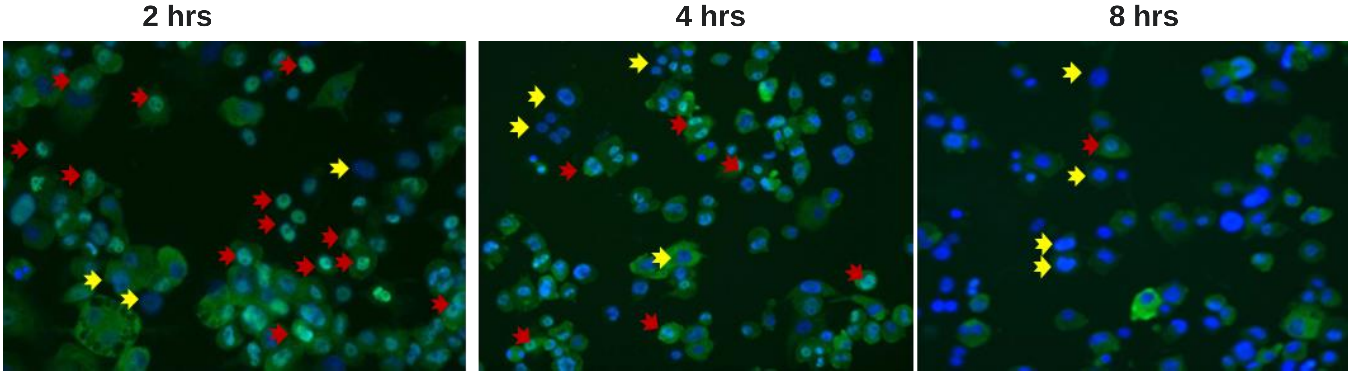

**Fig. S4. Nuclear translocation of pIRF3 following treatment of cells with SB 11285.** THP1 (WT) - derived macrophages were treated with S B 11285 alone, or DMSO control, for 2 hrs. Cells and the IRF3 translocation were monitored at 2, 4, and 8 hrs., post-treatment. Cells were fixed with 4% paraformaldehyde and treated with 0.5% Triton X-100, stained with rabbit anti-IRF3 and Alex Fluor 488 conjugated anti-rabbit IgG antibody (green). Nuclei were identified using DAPI (blue) staining. Cells were imaged on IXM (Molecular Devices) (40x). Images were analyzed using ImageJ and IRF3 and DAPI images were overlapped. Nuclear translocated p-IRF3 (green) is indicated by the red arrow. The yellow arrow (pointing to blue color labelled nucleus) indicates cells without IRF3 nuclear translocation. IRF3 nuclear translocation is a strong indicator that type I IFN signaling is activated. More red arrows means that more cells responded to the compound treatment. Magnification (40x).

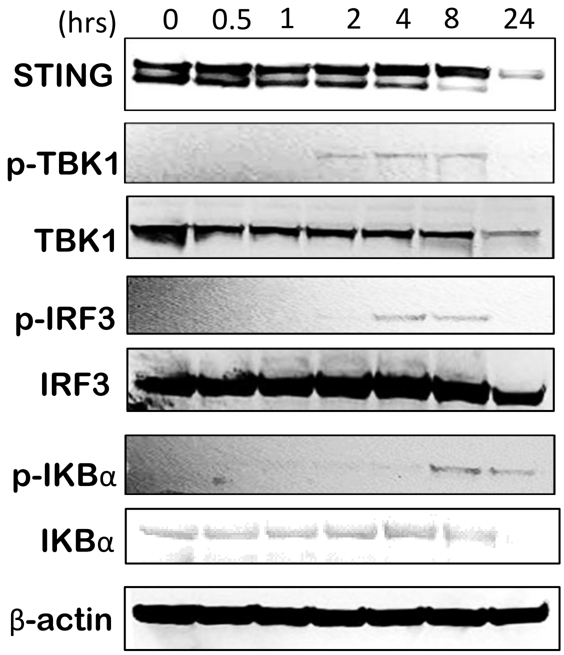

**Fig. S5. Characterization of Signaling Molecules in the STING Pathway After SB 11285 Treatment.** SB 11285 activation is characterized by signaling proteins indicated by formation of phosphorylated IRF3, TBK-1 and IκB-α, detectable after SB 11285 treatment. To ascertain this, THP-1 cells were treated with 5 µM SB 11285 with Lipofectamine LTX, and at different time-points, whole cell extracts were treated with RIPA lysis buffer and analyzed by immunoblotting with anti-hSTING antibody, anti-IRF3, anti-phospho-IRF3, anti-TBK1, anti-phospho TBK1, anti-IKBα, anti-phospho IKBα, β-actin antibodies and anti-rabbit IgG, HRP-linked antibody.

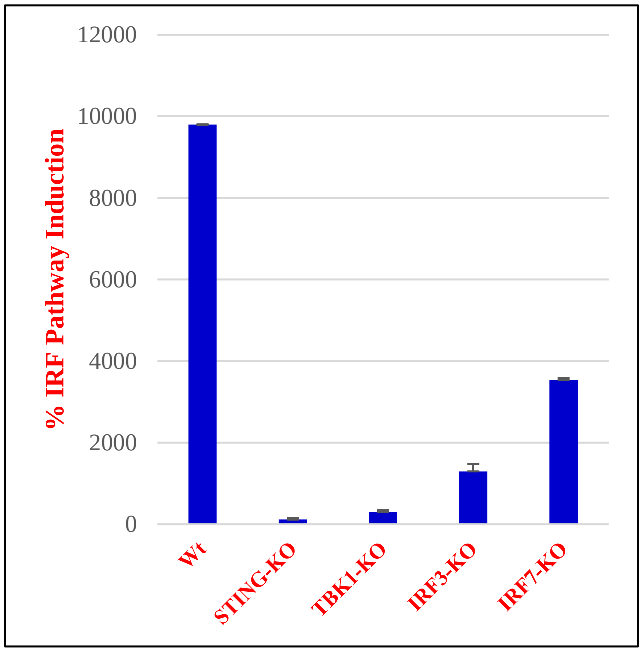

**Fig. S6. Characterization of Signaling Molecules in the STING Pathway After SB 11285 Treatment.** The role of STING, TBK1 & IRF3/7 in SB 11285-induced STING signaling was evaluated in Raw Macrophages. WT, STING KO, TBK1 KO, IRF3 KO and IRF7 KO RAW-Lucia ISG Cells were either treated with 50 µM of SB 11285 or DMSO. After 20 hrs., incubation, IRF activity was assessed using QUANTI-luc and the % induction was calculated from fold-change in luminescence/absorbance compared to DMSO-treated sample**.**

RAW-Lucia ISG cells, STING KO RAW-Lucia ISG cells, TBK1 KO RAW-Lucia ISG cells, IRF3 KO RAW-Lucia ISG cells, IRF7 KO RAW-Lucia ISG cells were used in this study. These cells carry Lucia gene under the control of an ISG54 minimal promoter to study IRF pathway. The cells grown in complete media were either treated with various concentrations of SB 11285 or DMSO control. After 20 hrs., incubation, the IRF activity was assessed using QUANTI-luc to measure the levels of Lucia. The % Induction was calculated from fold-change in luminescence/absorbance compared to the DMSO-treated sample.

**(A)** (B)

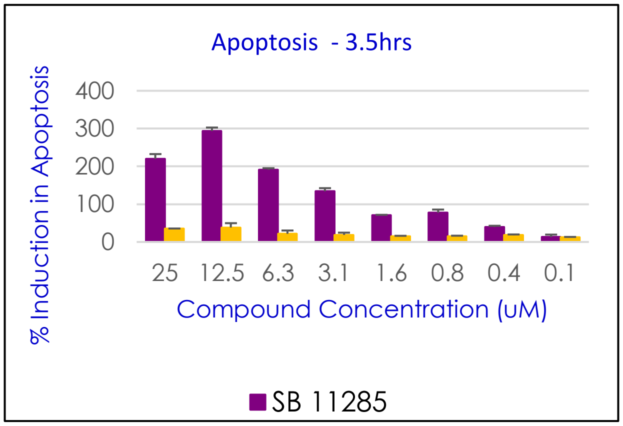

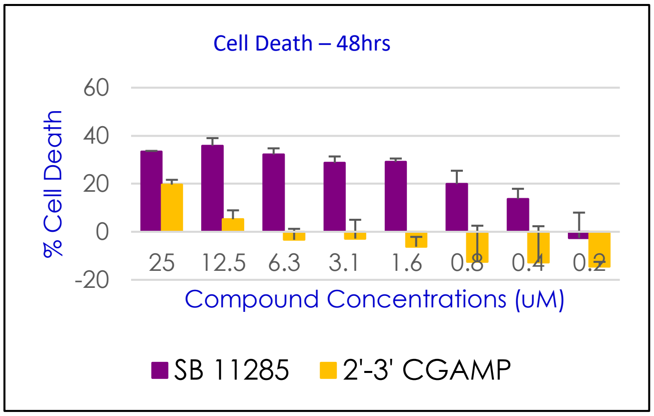

(C)

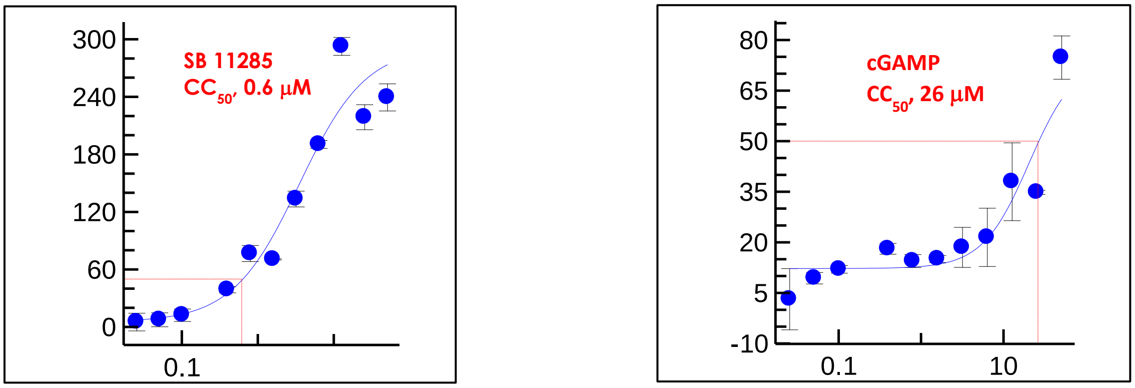

**Fig. S7. Evaluation of apoptosis and cell death in A20 mouse B cell Lymphoma cells. A.** A20 cells grown in complete media were treated with various concentrations of SB 11285, cGAMP or DMSO control with Lipofectamine LTX. After 4 hrs., incubation, % apoptosis was investigated by performing caspase 3/7 analysis using Caspase 3/7 Glo (Promega). **B**. After 48 hrs., incubation, the % cell death was assessed by performing cell survival analysis by Cell Titre Glo (Promega). **C**. The % cell death and/or apoptosis for SB 11285 and cGAMP was calculated from fold-change in luminescence compared to DMSO-treated sample. $CC_{50}$ values were generated by curve fit in Xlfit.

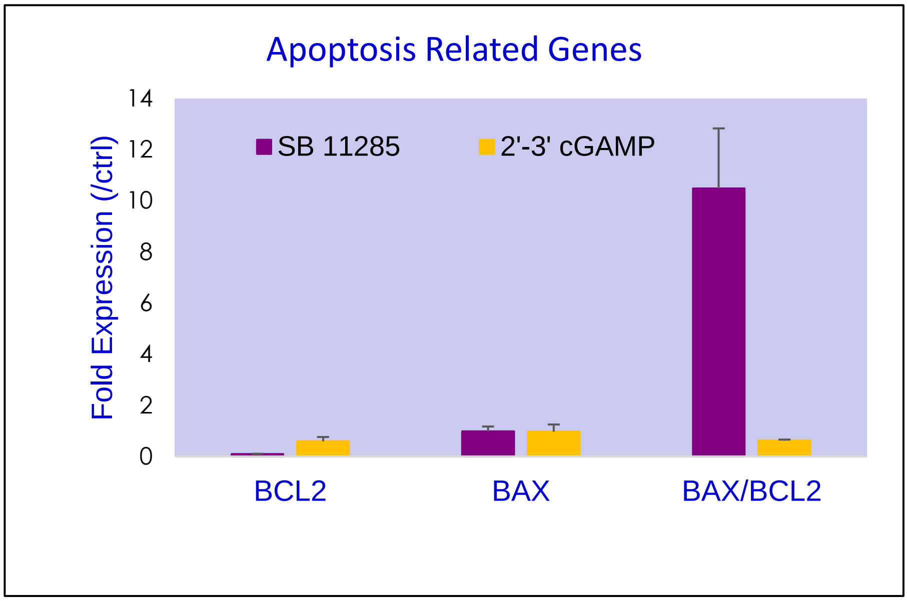

**Fig. S8. Gene expression analysis in A20 mouse B cell Lymphoma cells.** A20 cells grown in complete media were treated with 25 μM concentrations of SB 11285, cGAMP or DMSO control with Lipofectamine LTX. After 48 hrs., incubation, RNA was extracted, and gene expression of different genes related to apoptosis was evaluated by real time PCR. The fold-induction was calculated by ΔΔct method.

**(A)**

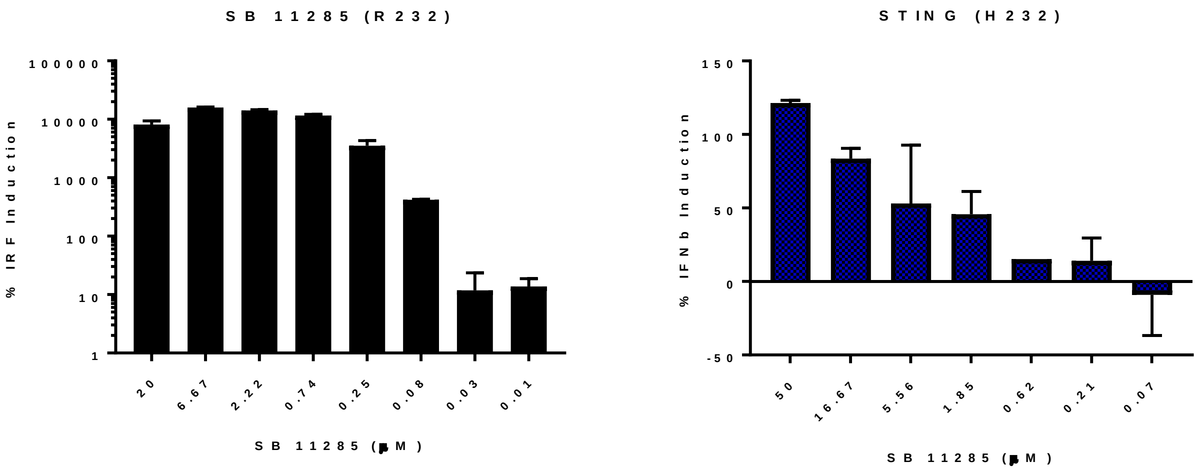

(B)

| Name | % Population | SB 11285 $EC_{50}$ (μM) | cGAMP $EC_{50}$ (μM) |
|---|---|---|---|
| hSTING [WT] | 57.9 | 0.008 | 3.9 |
| hSTING [HAQ] | 20.4 | 0.007 | 2.2 |
| hSTING [R232H] | 13.7 | 3.1 | ND |
| hSTING [AQ] | 5.2 | 0.43 | 25.4 |
| hSTING [R232Q] | 1.5 | 3.5 | 29.2 |
| mSTING | - | 0.15 | 3 |

**Fig. S9. Activity of SB 11285 Against STING Allelic Variants. (A).** To evaluate whether SB 11285 activates various allelic variants, THP1 cells naturally expressing HAQ isoform or STING-KO THP1 cells stably transfected with WT or allelic variants and carrying an ISRE-inducible Lucia and NF-κB inducible SEAP reporters were treated with various concentrations of SB 11285 or DMSO control. After 20 hrs., the levels of Lucia and SEAP were measured using QUANTI-luc and QUANTI-Blue to measure ISRE and NF-κB reporter activity respectively. The % IRF Induction was calculated from 5X fold-change in luminescence/absorbance in the treated samples compared to DMSO treated sample. $EC_{50}$ values were generated by curve fit in Xlfit. **(B).** Table showing $EC_{50}$ values for STING activation by SB 11285 of different STING alleles.

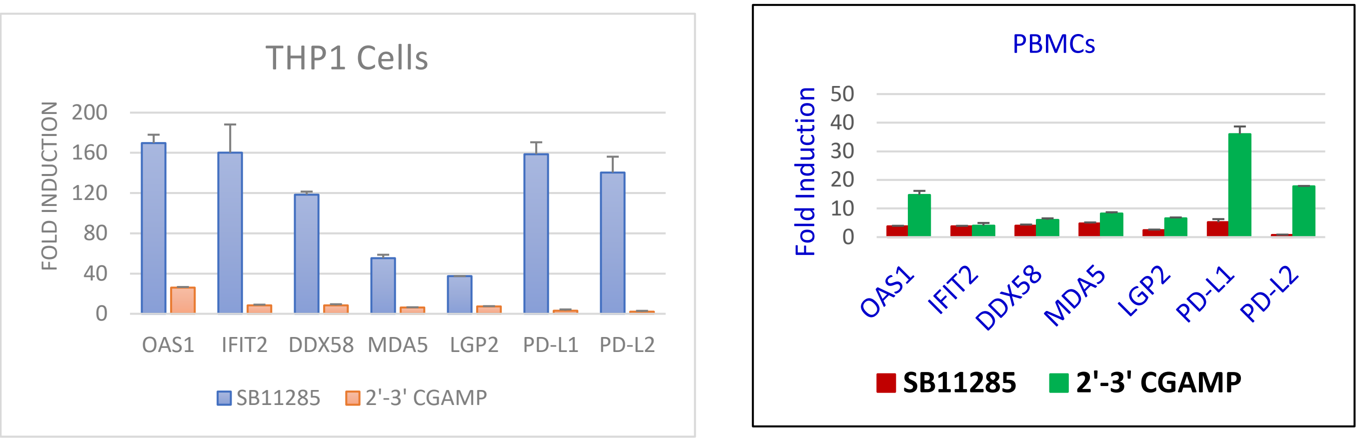

**Fig. S10. Gene Expression Analysis in THP1 and PBMCs.** The THP1 cells and PBMCs from healthy volunteers were grown in complete media were treated with 25 μM SB 11285 or cGAMP or DMSO with Lipofectamine LTX. After 48 hrs., incubation, RNA was extracted, and gene expression was evaluated by Real time PCR and fold-induction of genes was calculated by the ΔΔct method.

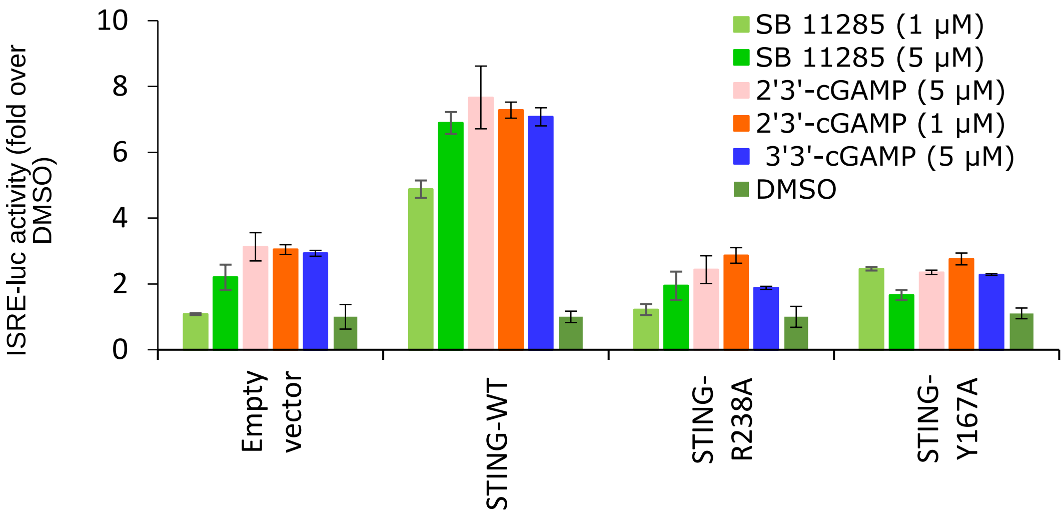

**Fig. S11. STING agonist activity of SB 11285 is abolished with STING mutants R238A and Y187A.** HEK293T cells were transfected with plasmids encoding wild-type STING or mutant STING carrying R238A or Y167A mutation together with ISG54 ISRE-luc reporter, as well as, renilla-luc internal reporter for 18 hrs., followed by treatment with 1 or 5 μM of SB 11285 in the presence of digitonin for 5 hrs. ISRE-luciferase activity was determined and normalized to Renilla luciferase internal control. Results are shown as fold-increase of normalized ISRE-luc activity over DMSO treated cells (mean ± standard deviation of triplicate wells per stimulant).

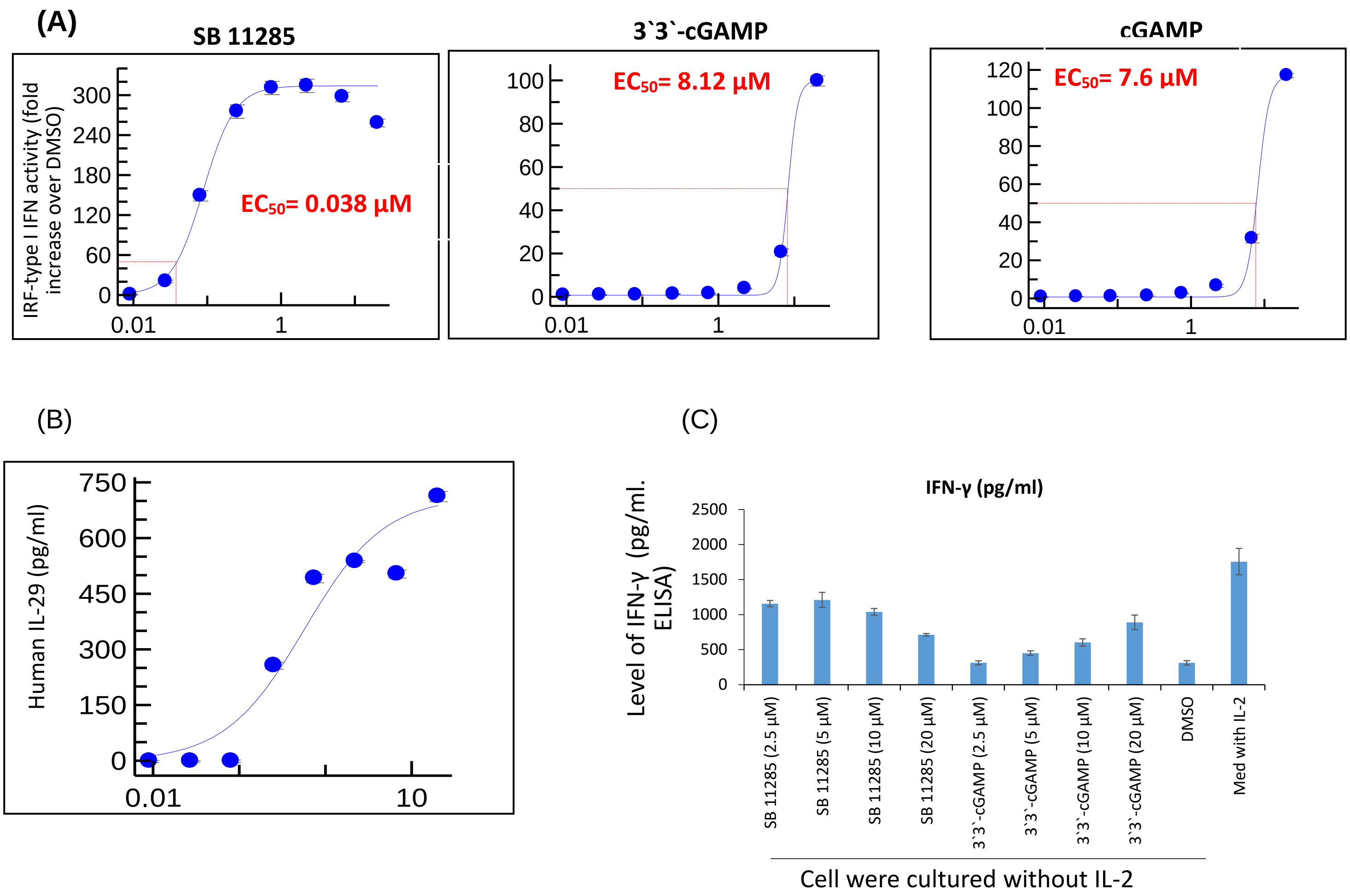

**Fig. S12. SB 11285 strongly activates both IRF-type I, Type II, and type III IFN responses.**

**A.** THP1-Dual cells were stimulated in triplicate with compound alone for 24 hrs. Levels of IRF-dependent secreted luciferase in cell culture supernatants for SB 11285, 3`,3`-cGAMP, and cGAMP treatments were measured using InvivoGen's Quanti-luc. Data shown are fold-induction over DMSO treated cells (mean ± standard deviation of triplicate wells per stimulant). **B**. Levels of IL-29 in culture supernatants were determined using ELISA. Results shown are average ± standard deviation of duplicate wells. Note: neither 3`,3`-cGAMP nor cGAMP induced detectable IL-29 in THP1 cells. **C**. NK92 cells were treated with SB 11285, 3`3`-cGAMP, or DMSO alone (without IL-2) for 21 hrs. Control cells were cultured in the presence of IL-2. Levels of IFN-γ in culture supernatants were measured using ELISA and is shown as pg/ml. (IL-2 can induce IFN-γ production in NK cells). Note: neither 3`,3`-cGAMP nor cGAMP induced detectable IFN-γ in the cells.

(A)

| Group | Number of mice | Dose | Dosing schedule | Route | MTV (day 12) mm³ |
|---|---|---|---|---|---|
| Vehicle | 10 | Saline | 1,4,7,10 & 17 | I.T. | 1196 |
| SB 11285 | 10 | 50 µg | 1,4,7,10 & 17 | I.T. | 222 |
| Anti-CTLA4 Ab | 10 | 5 mg/kg (followed by 2.5 mg/kg) | 1, (4 & 7) | I.P. | 906 |
| Anti-CTLA4 Ab + (SB 11285) | 10 | 5 mg/kg (followed by 2.5 mg/kg) + (50µg) | 1, (4 & 7) + (1,4,7,10 & 17) | I.P., + (I.T.) | 173 |

(B)

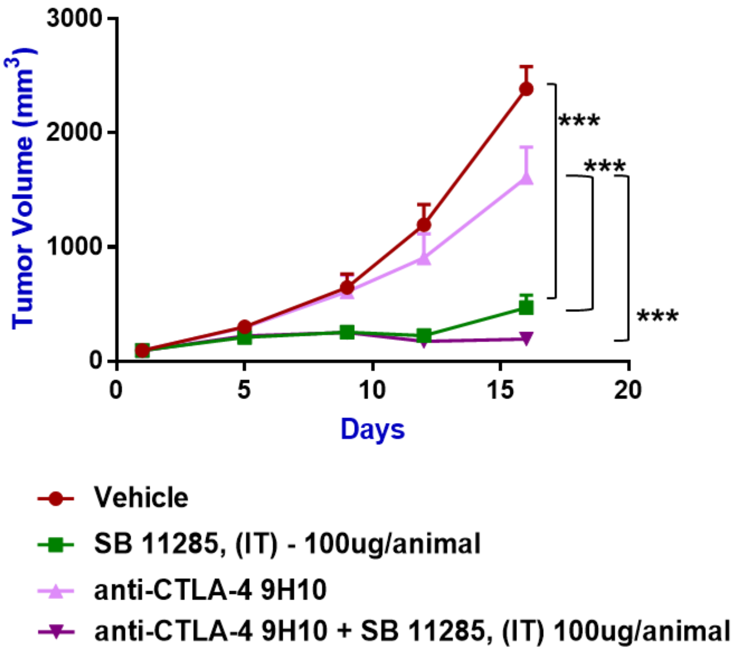

**Fig. S13**. **Anti-tumor activity of SB 11285 in combination with the anti-CTLA Antibody in the CT26 model**. **A**. Groups of female BALB/c mice (n = 10) were administered SB 11285 alone or in combination with anti-CTLA antibody as per the indicated schedule. The MTVs were recorded on a biweekly basis until MTV of 2000 $mm^3$ was reached in the vehicle group. **B**. MTV of each of the groups was plotted as shown. **Note.** The anti-tumor data is shown as MTVs measured on indicated days after initiation of treatment with the compounds. Additional experimental details and the criteria for statistical significance are given in the Materials and Methods Section.
